# *Plasmodium*-specific atypical memory B cells are not part of the long-lived memory response

**DOI:** 10.1101/350819

**Authors:** Damián Pérez-Mazliah, Peter J. Gardner, Edina Schweighoffer, Sarah McLaughlin, Caroline Hosking, Irene Tumwine, Randall S. Davis, Alexandre J. Potocnik, Victor Tybulewicz, Jean Langhorne

**Affiliations:** The Francis Crick Institute, London, UK; MRC National Institute for Medical Research, London UK; Departments of Medicine, Microbiology, and Biochemistry & Molecular Genetics, University of Alabama at Birmingham, Birmingham, AL, USA; School of Biological Sciences, The University of Edinburgh, Edinburgh, UK

## Abstract

A subset of atypical memory B cells accumulates in malaria and several infections, autoimmune disorders and aging in both humans and mice. It has been suggested these cells are exhausted long-lived memory B cells, and their accumulation may contribute to poor acquisition of long-lasting immunity to certain chronic infections, such as malaria and HIV. Here, we generated an immunoglobulin heavy chain knock-in mouse with a BCR that recognizes MSP1 of the rodent malaria parasite, *Plasmodium chabaudi*. In combination with a mosquito-initiated *P. chabaudi* infection, we show that *Plasmodium*-specific atypical memory B cells are short-lived and disappear upon natural resolution of chronic infection. These cells show features of activation, proliferation, DNA replication, and plasmablasts. Our data demonstrate that *Plasmodium*-specific atypical memory B cells are not a subset of long-lived memory B cells, but rather short-lived activated cells, and part of a physiologic ongoing B-cell response.

## INTRODUCTION

Atypical memory B cells (AMB) are an unusual B-cell subset detected in both mouse models and humans in the context of certain infections and autoimmune disorders, including HIV, HCV, tuberculosis, malaria, rheumatoid arthritis and systemic lupus erythematosus, and accumulated with age (Knox et al., 2017b; Naradikian et al., 2016a; Portugal et al., 2017; Rubtsov et al., 2017). In the context of infections, AMB were first described in HIV-viremic subjects, and termed tissuelike memory B cells, due to their similarity to an FCRL4-expressing memory B-cell subset found in human tonsillar tissues (Ehrhardt et al., 2005; Moir et al., 2008). In addition to FCRL4, these cells express relatively high levels of other potentially inhibitory receptors including CD22, CD85j, CD85k, LAIR-1, CD72, and PD-1, and show a profile of trafficking receptors including expression of CD11b, CD11c and CXCR3, consistent with migration to inflamed tissues. They are antigen-experienced class-switched B cells, which lack the expression of CD21 and the hallmark human memory B-cell marker CD27. Further studies demonstrated the expression of the transcription factor T-bet and the cytokine IFNγ by these cells, also characteristic of Th1 CD4+ T cells (Knox et al., 2017b; Obeng-Adjei et al., 2017; Portugal et al., 2017). Due to their poor functional capacity upon *in vitro* re-stimulation with BCR ligands, AMB were characterized as dysfunctional B cells, and increased frequencies of these cells was proposed to be a consequence of B-cell exhaustion driven by chronic inflammation and stimulation, drawing parallels with T-cell exhaustion during chronic viral infections (Moir et al., 2008; Portugal et al., 2015; Sullivan et al., 2015). It has been hypothesized that expansion of AMB might contribute to the mechanisms driving autoimmune disorders and deficiencies in acquisition of immunity to chronic infections. However, due to lack of good tools and animal models to analyze antigen-specific atypical B cells in greater depth, many of these concepts remain speculative.

Several studies suggest that AMB might contribute to poor acquisition of long-term immunity to *Plasmodium* infection (Illingworth et al., 2013; Portugal et al., 2015; Sullivan et al., 2015; 2016; Weiss et al., 2011; 2009; 2010). Indeed, some studies demonstrated that in the absence of constant re-exposure, *Plasmodium*-specific serum antibody levels rapidly wane, and full protection from clinical symptoms is lost, suggesting that B-cell memory is functionally impaired (Portugal et al., 2013). However, others have reported long-lasting maintenance of *Plasmodium*-specific antibodies and/or memory B cells in settings of differing malaria endemicity, and similar responses are also observed in mouse malaria models (Dorfman et al., 2005; Ndungu et al., 2009; 2013; Rasheed et al., 2012; Wipasa et al., 2010). Moreover, it has been shown that BCRs cloned from *P. falciparum*-specific AMB from malaria-exposed adults encode *P. falciparum*-specific IgG antibodies, which could contribute to *P. falciparum*-specific IgG antibodies in serum (Muellenbeck et al., 2013). These authors proposed that *P. falciparum*-specific AMB do not prevent, but rather contribute to the control of *Plasmodium* infection. These apparently contradictory results may reflect the fact that some studies were performed on the general peripheral blood B-cell pool and others focused on *Plasmodium*-specific B cells. In determining a role for these cells in a chronic infection it would be important to follow antigen-specific responses and to distinguish these from non-specific polyclonal B cell activation.

The study of the development of AMB is challenging and requires suitable mouse models, which allow for identification and isolation of antigen-specific B cells that exist often at very low frequency. Here, we generated a knock-in transgenic mouse with a high frequency of *Plasmodium chabaudi* Merozoite Surface Protein 1 (MSP1)_21_-specific B cells, to investigate memory B cells generated following mosquito-transmission of the rodent malaria, *P. chabaudi*. We identified a CD11b^+^CD11c^+^FCRL5^hi^ subset of MSP1_21_-specific B cells during the chronic infection with phenotypical and transcriptional features strikingly similar to those of human AMB. These AMB disappeared as the infection progressed, leaving a CD11b^−^CD11c^−^ FCRL5^h1^ MSP1_21_-specific B-cell compartment with characteristics of long-lived classical memory B cells (B_mem_) after the resolution of the infection. These shortlived MSP1 21-specific AMB were also generated in response to immunization, suggesting they may be a normal but transient component of a B-cell response to antigen. In this chronic *P. chabaudi* infection, it appears that AMB require ongoing antigenic stimulation driven by the sub-patent infection to persist, and do not represent a true long-lived “memory” B cell subset. Moreover, we show that generation of *Plasmodium*-specific AMB does not prevent the generation of *Plasmodium*-specific B_mem_, and does not prevent resolution of the infection.

## RESULTS

### Generation of an immunoglobulin heavy chain knock-in transgenic mouse model to study *Plasmodium*-specific B cell responses

To study Plasmodium-specific B cell responses in a rodent malaria model, we generated an /gh^NIMP23/+^ mouse strain on the C57BL/6J background (Materials and Methods and Figure S1).

The *Igh*^NIMP23/+^ mice were healthy, with no unusual behavioral or physical characteristics. There were no alterations in total cellularity, or in numbers of pro-B, pre-B, immature B, mature B and total B220^+^CD19^+^ B cells, plasma cells in the bone marrow of *Igh*^NIMP23/+^ mice (Figure 1A-B), as well as no alterations in the number of T1, T2, T3, follicular, marginal zone, and germinal center B cells, plasmablasts, plasma cells or total cellularity in the spleen of *Igh*^NIMP23/+^ mice (Figure 1C-D). Importantly, The *Igh*^NIMP23/+^ mice had a greatly increased frequency of B cells specific for MSP1_21_ (approximately 60% of the total B-cell compartment), as demonstrated by flow cytometry analysis of splenocytes with a MSP1_21_ fluorescent probe (Figure 1E-F).

**Figure 1.**
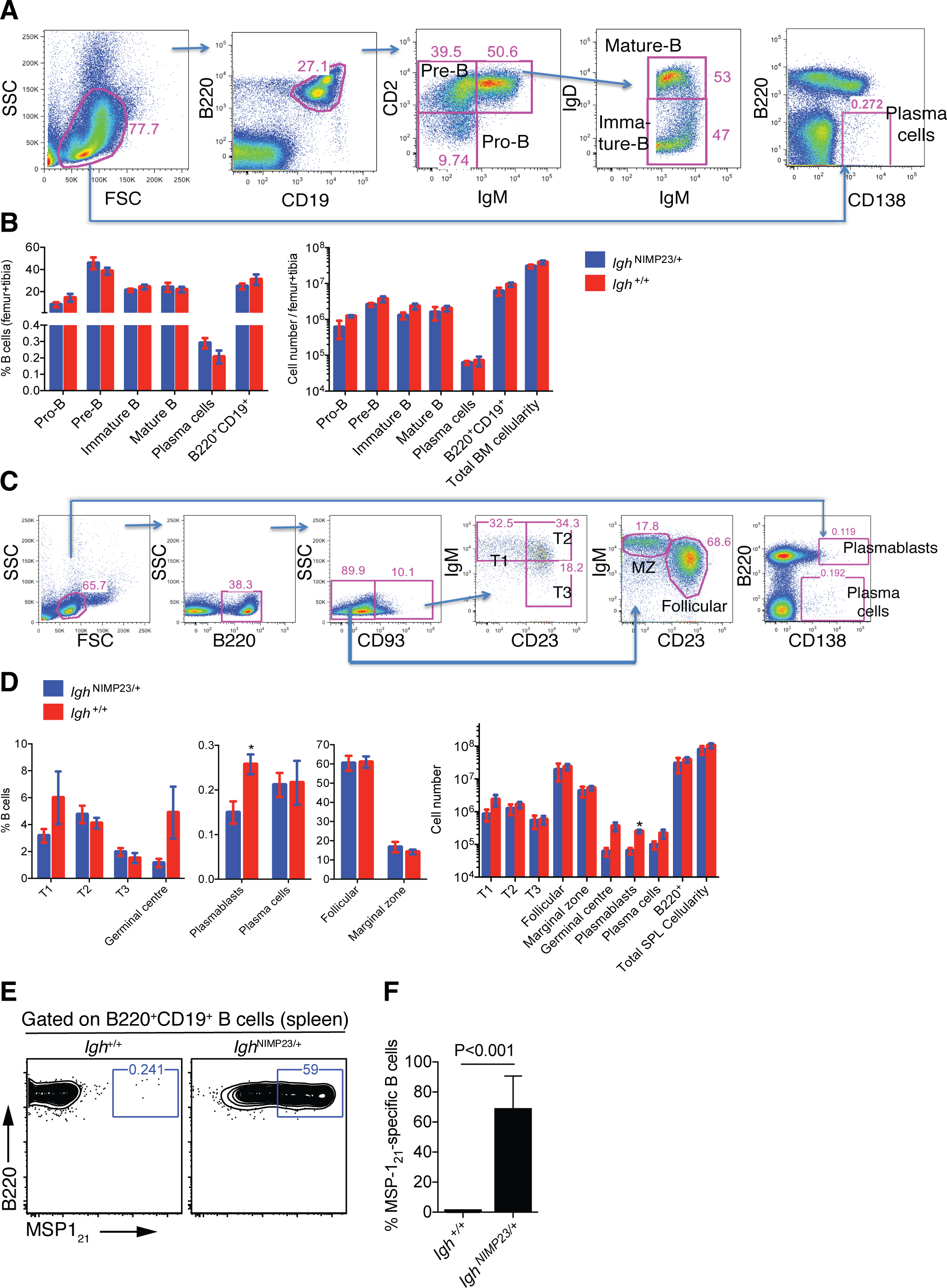
Analysis of total bone marrow and splenic B-cell populations in *Igh*^NIMP23/+^ and *Igh*^+^/^+^ littermates. (A) Flow cytometry gating strategy to identify different B-cell populations in bone marrow of *Igh*^NIMP23/+^ mice. Arrows indicate flow of analysis. The same strategy was used for *Igh*^+/+^ littermates. (B) Percentages and numbers of different B-cell populations in bone marrow of *Igh*^NIMP23/+^ and *Igh*^+/+^ littermates as defined in (A). (C) Flow cytometry gating strategy to identify different B-cell populations in spleen of *Igh*^NIMP23/+^ mice. (D) Percentages and numbers of different B-cell populations in spleen of *Igh*^NIMP23/+^ and *Igh*^+/+^ littermates as defined in (C). Data are representative of two independent experiments with 4 mice per group. (E) Flow cytometry analysis of B cells obtained from spleen of *Igh*^+/+^ (left) and *Igh*^NIMP23/+^ (right) mice stained with anti-B220 and CD19 antibodies in combination with an MSP1_21_ fluorescent probe. The gates show the frequency of B cells specific to MSP1_21_. (F) Frequencies of MSP1_21_-specific splenic B cells in *Igh*^NIMP23/+^ and wild-type *Igh*^+/+^ littermate controls (Mann Whitney U test). Data pooled from two independent experiments with 3-5 mice per group. Error bars are SEM. Mann Whitney U test.

### Increase in *Plasmodium*-specific B cells after mosquito transmission of *P. chabaudi*

To investigate B cells in *P. chabaudi* infections, which last several weeks, and to avoid potential problems with activation arising from very high frequencies of MSP1-specific B cells, we reduced the precursor frequency of MSP1_21_-specific B cells to match the natural level expected for antigen-specific B cells more closely, yet still readily detectable by flow cytometry. We generated mixed bone marrow (BM) chimeras by adoptively transferring a mixture of 10% bone marrow from either *Igh*^NIMP23/+^ or *Igh*^+/+^ mice (CD45.2+) together with 90% bone marrow from C57BL/6.SJL-*Ptprc*^a^ mice (CD45.1+) into sub-lethally irradiated *Rag2*^−/−^.C57BL/6.SJL-*Ptprc*^a^ mice (CD45.1+) to generate NIMP23→*Rag2*^−/−^ and WT→*Rag2*^−/−^ bone marrow chimeric mice respectively (Figure S2A-B). In both types of chimeras, 2-3% of the B cells were CD45.2^+^ and in NIMP23→*Rag2*^−/−^ mice approximately 1-2% of the B cells were MSP1_21_-specific (Figure S2C-E). No MSP1_21_-specific B cells were detected in the control WT→*Rag2*^−/−^ chimeras (Figure S2D).

Infection of C57Bl/6J *wt* mice with *P. chabaudi* by mosquito bite gives rise to a short (48h) pre-erythrocytic infection, followed by an acute blood parasitemia peaking approximately 10d post-transmission. Thereafter, the infection is rapidly controlled, reaching very low parasitemias by 15d post-transmission, with a subsequent prolonged (~90d), but low-level chronic infection before parasite elimination (Brugat et al., 2017; Spence et al., 2013). NIMP23→*Rag2*^−/−^ mice infected with *P. chabaudi* by mosquito bite, showed a similar course of parasitemia to that of control WT→*Rag2*^−/−^ mice (Figure S2F), and C57BL/6J *wt* mice (Brugat et al., 2017; Spence et al., 2013; 2012). Importantly, the MSP1_21_-specific *Igh*^NIMP23/+^ B cells (CD45.2^+^MSP1_21_^+^) in NIMP23→*Rag2*^−/−^ chimeras showed a robust response to the infection, as demonstrated by a dramatic increase in the proportions and numbers of GL-7^+^CD38^lo^ germinal centers (GC) and IgG2b+IgD^−^ class-switched B cells in the spleen at 35 days post-infection (dpi) (Figure S2G-H).

Thus, we have generated a mouse model with a detectable numbers of functional MSP1_21_-specific B cells capable of responding to *P. chabaudi* infection and which can be readily followed.

### Generation of *Plasmodium*-specific AMB after mosquito transmission of *P. chabaudi* infection

We investigated whether *Plasmodium*-specific AMB could be identified in mice during a blood-stage *P. chabaudi* infection. We selected a series of mouse homologues to human cell surface markers described on human AMB (Charles et al., 2011; Kardava et al., 2014; 2011; Knox et al., 2017a; H. Li et al., 2016; Moir et al., 2008; Muellenbeck et al., 2013; Portugal et al., 2015; Russell Knode et al., 2017; Sullivan et al., 2015). Human AMB express CD11b, CD11c, Fc receptor-like (FCRL) 3-5, high levels of CD80, low levels of CD21, and are Ig class-switched. Mouse FCRL5 most closely resembles human FCRL3 and is the only mouse FCRL-family member which contains both ITIM and ITAM motifs in its cytoplasmic tail (Davis, 2007; 2004; Won et al., 2006; Zhu et al., 2013). Therefore, our flow cytometry panel for mouse AMB included antibodies against CD11b, CD11c, FCRL5, CD21, IgD, and also CD80 and CD273 which identify mouse B cells that are antigen-experienced and potentially memory cells (Anderson et al., 2007; Tomayko et al., 2010; Zuccarino-Catania et al., 2014).

We detected an increased number of cells in a distinct CD11b^+^CD11c^+^ MSP1_21_-specific B-cell subset at 28-35dpi, in the chronic phase of *P. chabaudi* infection (Figure 2A-B). This subset showed several AMB characteristics, including high expression of FCRL5 and low expression of CD21 and IgD (Figure 2C-E). In addition, the CD11b^+^CD11c^+^ MSP1_21_-specific B-cell subset was enriched with cells expressing CD80 and CD273 (Figure 2C-E).

We then explored whether this CD11 b^+^CD11c^+^ MSP1_21_-specific B cell subset was detected during the memory phase, i.e. after resolution of the infection. As it takes up to 90 days for a blood-stage *P. chabaudi* infection to be eliminated from C57BL/6J mice (Achtman et al., 2007; Spence et al., 2013), we measured these responses from 155dpi onwards. Unexpectedly, the numbers of CD11b^+^CD11c^+^ MSP1_21_-specific B cells were not significantly higher than background level (Figure 2A-B).

These data demonstrate that a mosquito-borne infection with *P. chabaudi* generates *Plasmodium*-specific B cells resembling human AMB. However, these cells do not persist and are not detected above background level after parasite clearance.

**Figure 2.**
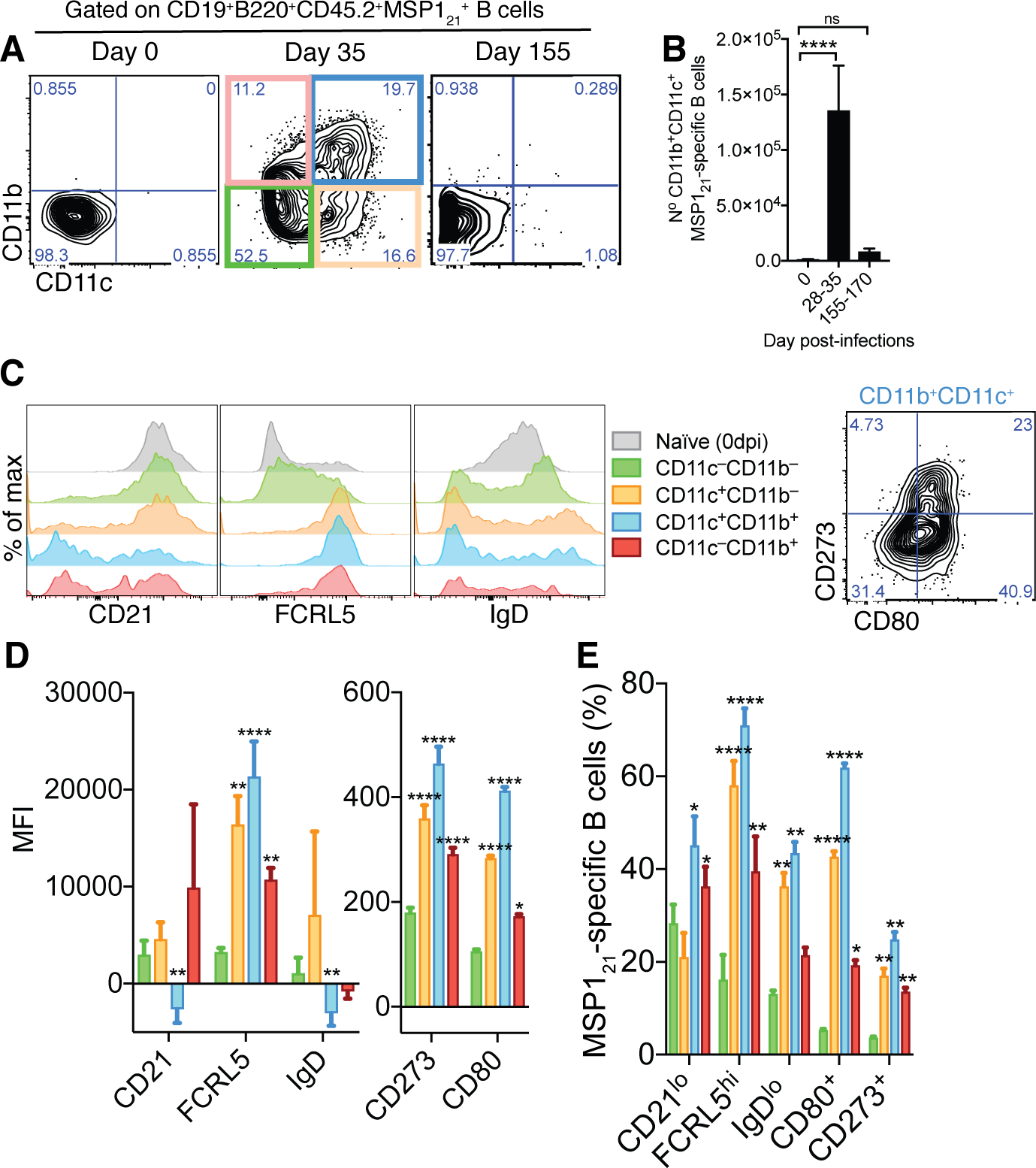
Generation of MSP1_21_-specific AMB in response to mosquito transmitted *P. chabaudi* infection. (A) Flow cytometry showing differential expression of CD11b and CD11c on splenic MSP1_21_-specific B cells from NIMP23→*Rag2*^−/−^ chimeric mice before infection (day 0) and at 35 and 155dpi. (B) Numbers of splenic MSP1_21_-specific CD11b^+^CD11c^+^ AMB from NIMP23→*Rag2*^−/−^ during the course of mosquito transmitted *P. chabaudi* infection. Kruskal-Wallis test vs day 0. ****, P<0.0001 (C) Flow cytometry showing expression of CD21/35, FCRL5, IgD, CD273 and CD80 on different subsets of splenic MSP1_21_-specific B cells from NIMP23→*Rag2*^−/−^ chimeric mice defined based on CD11b and CD11c expression at 35dpi. (D) Geometric mean fluorescence intensity (MFI) of CD21/35, FCRL5, IgD, CD273 and CD80 expression on different subsets of splenic MSP1_21_-specific B cells from NIMP23→*Rag2*^−/−^ chimeric mice defined based on CD11b and CD11c expression at 35dpi. (E) Frequencies of CD21/35, FCRL5, IgD, CD273 and CD80 positive cells among different subsets of splenic MSP1_21_-specific B cells from NIMP23→*Rag2*^−/−^ chimeric mice defined based on CD11b and CD11c expression at 35dpi. Two-way ANOVA vs CD11b^−^CD11c^−^ subset. *, P<0.05; **, P<0.01, ***, P<0.001; ****, P<0.0001. Error bars are SEM. Data pooled from three independent experiments with 3-5 mice per group.

### Transcriptome analysis confirms the AMB nature of CD11b^+^CD11c^+^ MSP1_21_-specific B cells, and reveals a plasmablast-like signature for this subset

To gain a better understanding of the identity of the CD11b^+^CD11c^+^ *Plasmodium*-specific B cell subset, we isolated both CD11b^+^CD11c^+^ and CD11b^−^CD11c^−^ MSP1_21_-specific B cells from spleens of *P. chabaudi*-infected NIMP23→*Rag2*^−/−^ mice (35dpi) (Figure S3), and MSP1 21-specific B cells from the spleen of naive NIMP23→*Rag2*^−/−^ mice (Figure 2A), by flow cytometric sorting, and performed an mRNAseq transcriptional analysis on the three populations.

The transcriptome of CD11b^+^CD11c^+^ MSP1_21_-specific B cells highly resembled that of human AMB. A series of hallmark genes upregulated in human AMB were also upregulated in CD11b^+^CD11c^+^ MSP1 21-specific B cells, including IgG (*Ighg2c* and *Ighg2b*), *Cxcr3*, *Tbx21* (T-bet), *Lair1* and *Fcrl5* (Figure 3, genes in red boxes, and references in Table S1). In addition, the MSP1_21_-specific CD11b^+^CD11c^+^ B-cell subset showed upregulation of *Ifng*, *Aicda*, a large array of inhibitory receptors [including *Pd1*, *Cd72*, *Cd85k*, *Fcgr2b* (CD32b), *Siglece*], antigen-experienced/memory markers [*Cd80*, *Cd86*, *Nt5e* (CD73) and high *Cd38*], and additional class-switched immunoglobulins (i.e. *Igha*, *Ighg1* and *Ighg3*), all of which have been shown to be upregulated on human AMB (Figure 3C-G, references in Table S1). These cells also expressed Galectins (*Lgals1* and *Lgals3*), previously implicated in B-cell anergy (Clark et al., 2007) (Figure 3H), and displayed a pro-apoptotic program (e.g. high expression of *Fasl*, and low expression of *Bcl2*) (Figure 3I). Interestingly, in agreement with data on human AMB, MSP1_21_-specific CD11b^+^CD11c^+^ B cells showed upregulation of *Mki67* (Figure 3J), indicative of proliferation, and had characteristics of plasmablasts and/or plasma cells, including upregulation of *Cd138*, *Prdm1* (*Blimp1*) and *Xbp1*, and low expression of *Cxcr5*, *Pax5*, and *Bcl6* (Figure 3B and K). However, these cells showed low expression of *Irf4* and *S1p1*, suggesting that they may be in a pre-plasmablast or pre-migratory plasma-cell stage (Kabashima et al., 2006; Kallies et al., 2007) (Figure 3K). Finally, and similar to human AMB, CD11b^+^CD11c^+^ MSP1_21_-specific B cells showed low expression of *Cd40*, *Cr2* (CD21), *Ms4a1* (CD20), and *Cd24a* (Figure 3L).

**Figure 3.**
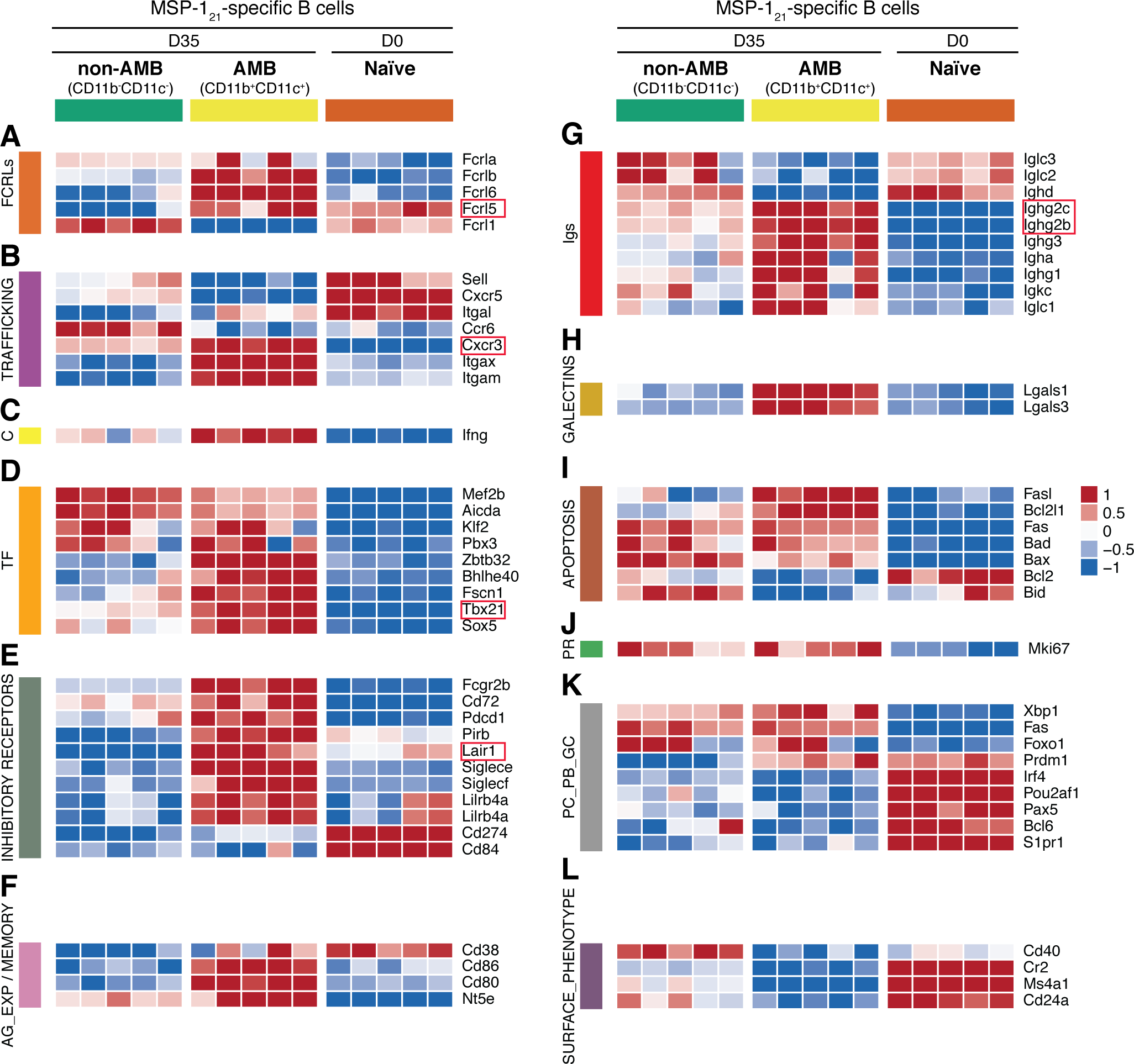
Transcriptome analysis of sorted splenic MSP1_21_-specific CD11b^+^CD11c^+^ AMB. MSP121-specific CD11b^+^CD11c^+^ (AMB) and CD11b“CD11c” B cells were flow cytometry sorted from the spleen of NIMP23→*Rag2*^−/−^ chimeric mice at 35dpi; MSP1 21-specific B cells were flow cytometry sorted from the spleen of naive NIMP23→*Rag2*^−/−^, and these three B cell populations were submitted to mRNAseq analysis. The heat maps display level of expression of selected individual genes, organized in functional clusters related to (A) Fc receptor like molecules, (B) cell trafficking, (C) cytokines, (D) transcription factors, (E) inhibitory receptors, (F) antigen experience/memory, (G) immunoglobulins, (H) galectins, (I) apoptosis, (J) proliferation, (K) plasma cells/plasmablasts/germinal centers, (L) surface markers. Each column corresponds to data from an individual mouse (n=5 35dpi, n=5 0dpi).

We ran a Gene Set Enrichment Analysis (GSEA) (Subramanian et al., 2005) with a gene list ranked according to their differential expression between MSP1_21_-specific CD11b^+^CD11c^+^ AMB sorted from infected mice and MSP1_21_-specific B cells sorted from naïve mice, using gene sets *a priori* obtained from Reactome (Fabregat et al., 2017). Among the gene sets yielding the top 50 significant (fdr<0.001) highest normalized enrichment score (*NES*) we obtained gene sets corresponding to cell cycle, DNA replication, generation/consumption of energy, regulation of apoptosis, activation of NF-ᴋB on B cells, and downstream signaling events of the BCR (Table S2 and Figure S4). These data further corroborate the activated and proliferative nature of MSP1_21_-specific CD11b^+^CD11c^+^ AMB.

Taken together, these data demonstrate that CD11b^+^CD11c^+^ MSP1_21_-specific mouse AMB present during the chronic phase of *P. chabaudi* infection are very similar to human AMB described in several chronic infections. In addition, this B-cell subset shows features of activation, proliferation, DNA replication and plasmablasts, resembling previous observations in human AMB (Muellenbeck et al., 2013).

### Generation of *Plasmodium*-specific AMB in response to immunization

The occurrence of CD11b^+^CD11c^+^ AMB might be a consequence of aberrant B-cell activation driven exclusively by certain pathogens. Alternatively, they might be part of a normal B-cell response, which is exacerbated by the persistent nature of certain infections. To test whether CD11b^+^CD11c^+^FCRL5^+^ AMB could be generated in the absence of persistent infection, we immunized mice with MSP1_21_. A previous report had demonstrated the presence of CD11b^+^CD11c^+^Tbet^+^ B-cells 24h postimmunization with R848, a TLR7/8 ligand (Rubtsova et al., 2013). Therefore, we immunized *Igh*^NIMP23/+^ mice with R848 together with the antigen MSP1_21_ and looked for the appearance of MSP1_21_-specific CD11b^+^CD11c^+^FCRL5^+^ atypical B cells. We observed substantial numbers of MSP1_21_-specific CD11b^+^CD11c^+^ B cells in the spleens of *Igh*^NIMP23/+^ mice 24h post-immunization (Figure 4A). These cells expressed increased levels of both FCRL5 and CD80 (Figure 4B-C) and did not display GC characteristics (Figure 4D), similar to the MSP1_21_-specific CD11b^+^CD11c^+^FCRL5^+^ atypical B cells generated following *Plasmodium* infection. The MSP1_21_-specific CD11b^+^CD11c^+^ B cells observed after immunization appeared only transiently, as they could no longer be detected at 3 and 7d post-immunization (Figure 4A and E).

These data demonstrate that MSP1_21_-specific CD11b^+^CD11c^+^ AMB can be generated independently of the infection, and that they are short-lived cells.

**Figure 4.**
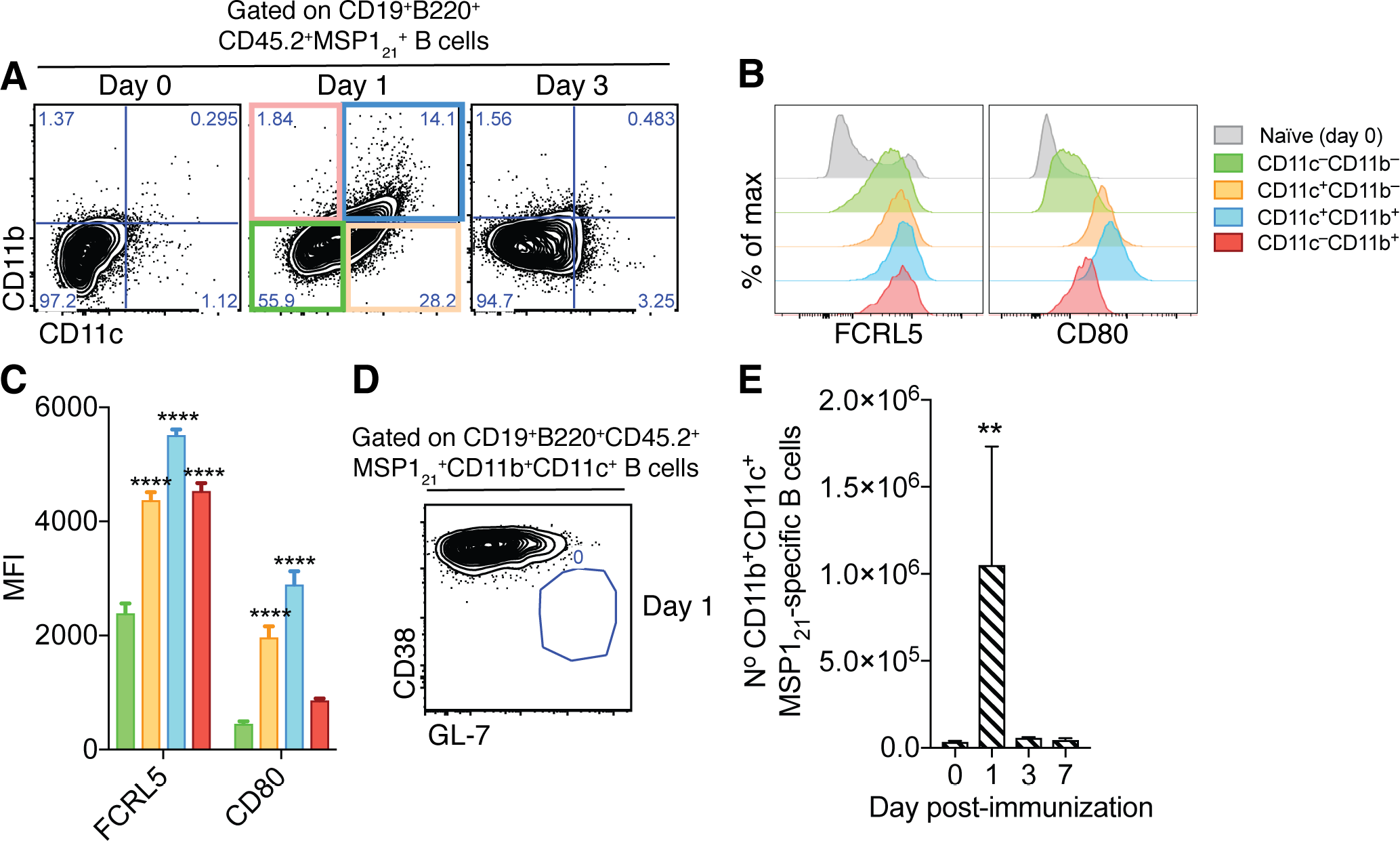
Generation of splenic MSP1_21_-specific CD11b^+^CD11c^+^ AMB in response to immunization. (A) Flow cytometry showing differential expression of CD11b and CD11c on splenic MSP1_21_-specific B cells from *Igh*^NIMP23/+^ mice before immunization (day 0) and at days 1 and 3 post-immunization with R848 and MSP1_21_. (B) Flow cytometry showing expression of FCRL5 and CD80 on different subsets of splenic MSP1_21_-specific B cells from *Igh*^NIMP23/+^ defined based on CD11b and CD11c expression at day 1 post-immunization and naive mice. (C) Geometric MFIof FCRL5 and CD80 expression on different subsets of splenic MSP1_21_-specific B cells from *Igh*^NIMP23/+^ defined based on CD11b and CD11c expression at day 1 post-immunization. Two-way ANOVA vs CD11b^−^CD11c^−^ subset. ****, P<0.0001. (D) Flow cytometry of CD38 vs GL-7 (GC markers) on CD11b^+^CD11c^+^ MSP1_21_-specific B cells from Igh^NIMP23/^+ at day 1 post-immunization. (E) Numbers of splenic CD11b^+^CD11c^+^ MSP1_21_-specific B cells from *Igh*^NIMP23/+^ during the course of immunization. Kruskal-Wallis test compared to day 0. **, P<0.01. Error bars are SEM. Data pooled from three independent experiments with 3-5 mice per group.

### *Plasmodium*-specific CD80^+^CD273^+^ B_mem_ are generated and persist after resolution of *P. chabaudi* infection

Identification of mouse B_mem_ by flow cytometry originally relied on detecting B cells that had undergone Ig class-switching from IgM to IgG, and that did not express GC markers (i.e. IgG^+^CD38^hi^GL-7^lo^) (Lalor et al., 1992; Ridderstad and Tarlinton, 1998). More recently, this set of markers has been extended to include CD80, CD273 (PD-L2) and CD73, with CD273 and CD80 being the most useful to discriminate memory from naïve B cells (Tomayko et al., 2010; Zuccarino-Catania et al., 2014). In combination, these markers allow the identification of different subsets of switched as well as non-class switched (i.e. IgM/D^+^) B_mem_. Therefore, we used cell surface expression of CD80 and CD273 on MSP1_21_-specific B cells to identify B_mem_ during and after resolution of *P. chabaudi* infection.

MSP1_21_-specific B cells from spleens of naive NIMP23→*Rag2*^−/−^ mixed BM chimeras showed little expression of either CD80 or CD273 (Figure 5A). By contrast, CD80^+^CD273^+^, CD80^+^CD273^−^ and CD80^−^CD273^+^ MSP121-specific B cells, both class-switched (IgM/D^lo^) and non-class-switched (IgM/D^hi^), were readily detected above background levels at 28-35dpi in *P. chabaudi*-infected NIMP23→*Rag2*^−/−^ mixed BM chimeras (Figure 5B and D). After resolution of infection (155-170dpi), the numbers of CD80^+^CD273^+^ and CD80^+^CD273^−^ MSP1_21_-specific B cells were either sustained or increased, while the CD80^−^CD273^+^ population decreased, compared with 28-35dpi (Figure 5A-D). All three MSP1 21-specific Bmem subsets showed high CD38 expression (Figure 5E, F). Importantly, the MSP1_21_-specific GC B cells detected at 28-35dpi did not express either CD80 or CD273, further distinguishing GC from memory and atypical memory *P. chabaudi*-specific B-cell subsets (Figure 5G). No MSP1_21_-specific GC B cells were detected above background level after resolution of the infection (Figure 5H). Finally, and in accordance with Figure 3, approximately 70% of the MSP1_21_-specific CD11b^+^CD11c^+^ AMB observed at days 28-35pi expressed either CD80, CD273 or a combination of both (Figure 5I).

These data show that, in contrast to the transient AMB, splenic CD80^+^CD273^+^ and CD80^+^CD273^−^ class-switched and non-class-switched MSP1_21_-specific B_mem_ persist after resolution of *P. chabaudi* infection.

**Figure 5.**
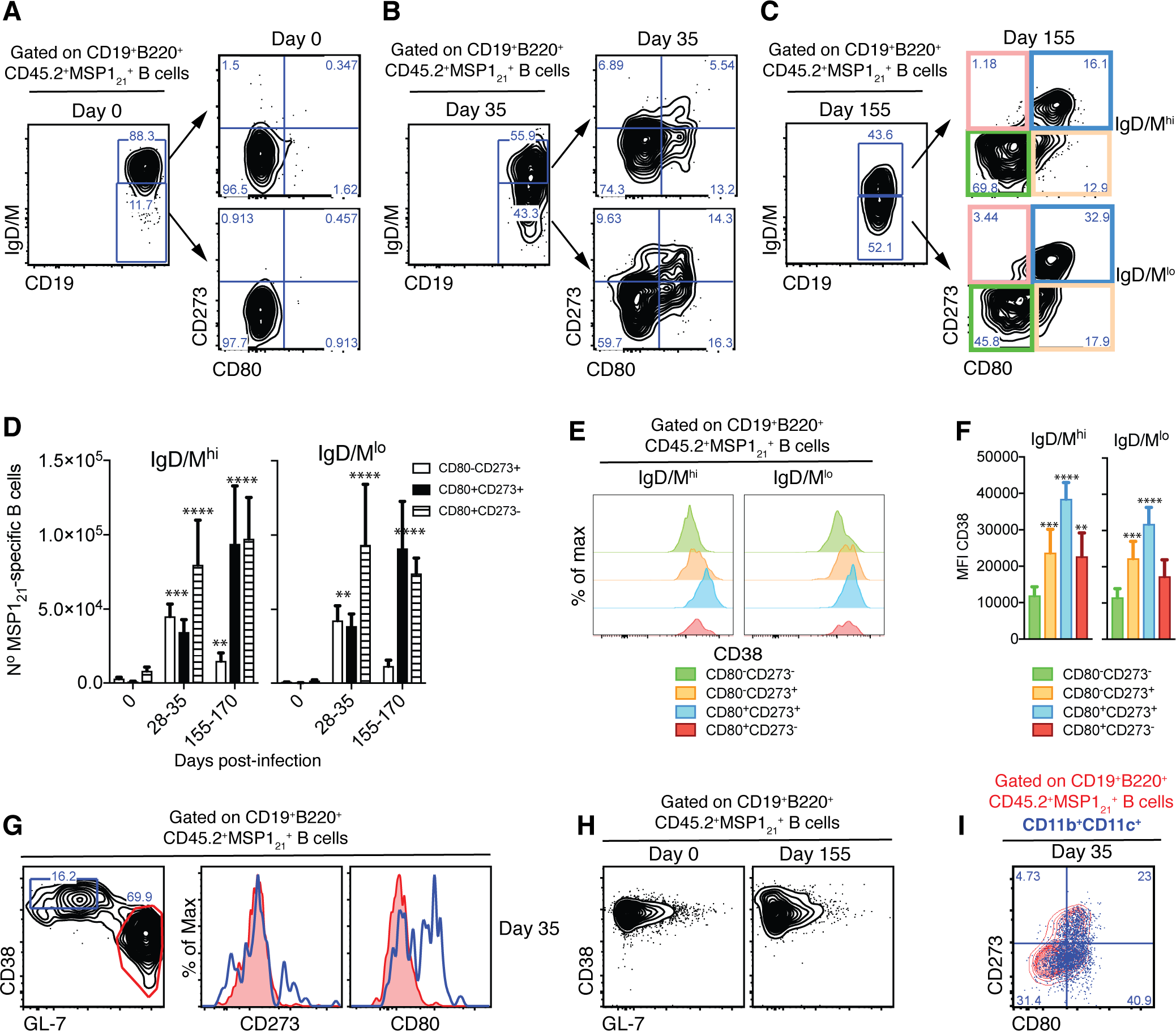
Detection of MSP1_21_-specific B_mem_ after resolution of *P. chabaudi* infection. (A), (B) and (C) Flow cytometry showing gating strategy to identify splenic IgM/D^hi^ and IgM/D^lo^ CD273^+^ and/or CD80^+^ MSP1_21_-specific Bmem in NIMP23→*Rag2*^−/−^ chimeric mice before infection (day 0), at 35 and 155dpi, respectively. (D) Numbers of splenic IgM/D^hi^ and IgM/D^lo^ CD273^+^ and/or CD80^+^ MSP1_21_-specific B_mem_ in NIMP23→*Rag2*^−/−^ chimeric mice during the course of mosquito transmitted *P. chabaudi* infection. Two-way ANOVA vs day 0. **, P<0.01; ***, P<0.001; ****, P<0.0001. (E) Flow cytometry showing expression of CD38 on different subsets of splenic IgM/D^hi^ and IgM/Dμ MSP1 21-specific B cells from NIMP23→*Rag2*^−/−^ chimeric mice defined based on CD273 and CD80 expression at 155dpi. (F) Geometric MFIof CD38 expression on different subsets of IgM/D^hi^ and IgM/D^lo^ splenic MSP1_21_-specific B cells from NIMP23→*Rag2*^−/−^ chimeric mice defined based on CD273 and CD80 expression at day 155 post-mosquito transmitted *P. chabaudi* infection. Two-way ANOVA vs CD273^−^ CD80^−^ subset. **, P<0.01; ***, P<0.001; ****, P<0.0001. Error bars are SEM. (G) Flow cytometry of CD273 and CD80 expression on non-GC (CD38^hi^GL-7^lo^, blue) and GC (CD38^lo^GL-7^h1^, red) splenic MSP1_21_-specific B cells from NIMP23→*Rag2*′^−/−^ chimeric mice at 35dpi. (H) Flow cytometry of CD38 vs GL-7 (GC markers) on splenic MSP1_21_-specific B cells from NIMP23→*Rag2*^−/−^ chimeric mice at 0 and 155dpi. (I) MSP1_21_-specific B cells CD11b^+^CD11c^+^ MSP1_21_-specific B cells overlaid on the CD80 vs CD273 plot corresponding to total MSP1_21_-specific B cells. Data pooled from three independent experiments with 3-7 mice per group.

### *Plasmodium*-specific B_mem_ express high levels of FCRL5

As discussed above, no single marker has been described so far that can identify all mouse B_mem_ subsets. Surprisingly, we observed that after resolution of infection (155-170dpi), MSP1_21_-specific B cells expressing different combinations of CD80 and CD273 (CD80^+^CD273^+^, CD80^−^CD273^+^ or CD80^+^CD273^−^) all expressed very high levels of FCRL5, in contrast to CD80^−^CD273^−^ MSP1_21_-specific B cells that express no memory markers at this stage (Figure 6A). This suggests that FCRL5 might be a marker for all antigen-experienced B cells. In order to confirm this, we used unsupervised methods to analyze our multiparameter flow cytometry data. We used PhenoGraph and t-SNE within the Cytofkit package (Materials and Methods, Chen et al 2016) to analyze MSP1_21_-specific B cells based on the expression of FCRL5, CD38, IgD, CD80 and CD273 on these cells, as determined by flow cytometry (Figure 6A and B). The analysis identified six clusters of cells with memory characteristics displaying high expression of CD38, CD80 and/or CD273, and variable expression of IgD, all of which expressed high levels of FCRL5 (Figure 6C: clusters identified with purple arrows). We then used Isomap, (Cytofkit package) to infer the relatedness between those cell subsets identified by PhenoGraph. This confirmed high similarities between the cell clusters expressing high levels of FCRL5 with the clusters expressing high levels of the memory markers CD80, CD273 and CD38 (Figure 6D).

To confirm the memory identity of MSP1_21_-specific FCRL5^hi^ B-cells detected after resolution of the infection, we isolated MSP1_21_-specific B cells expressing either high levels of FCRL5 or not expressing FCRL5 (i.e. FCRL5^hi^ and FCRL5^−^ MSP1_21_-specific B cells) from the spleen of *P. chabaudi*-infected NIMP23→*Rag2*^−/−^ mice (155dpi) (Figure S5), and MSP1 21-specific B cells from the spleen of naive NIMP23→*Rag2*^−/−^ mice, by flow cytometric sorting, and performed mRNAseq analysis on these three sorted cell populations (Figure 6E). As expected, the MSP1_21_-specific FCRL5^hi^ B cell subset showed high expression of genes encoding the hallmark memory B cell markers *Cd38*, *Cd80*, *Cd86*, *Nt5e* (CD73) and *Pdcd1lg2* (CD273), when compared with either MSP1_21_-specific FCRL5− B cells sorted at the same time or MSP1_21_-specific B cells sorted from naïve mice (Figure 6E). Moreover, the MSP1_21_-specific FCRL5^hi^ B cells sorted after resolution of the infection upregulated the anti-apoptotic *Bcl2* gene, which is an additional hallmark characteristic of memory B cells (Figure 6E). Importantly, FCRL5 also identified CD80^+^ and CD273^+^ MSP1_21_-specific B_mem_ subsets generated following immunization with a model antigen (Figure S6).

Thus, after resolution of the infection, high expression of FCRL5 identifies *P. chabaudi*-specific B_mem_.

**Figure 6.**
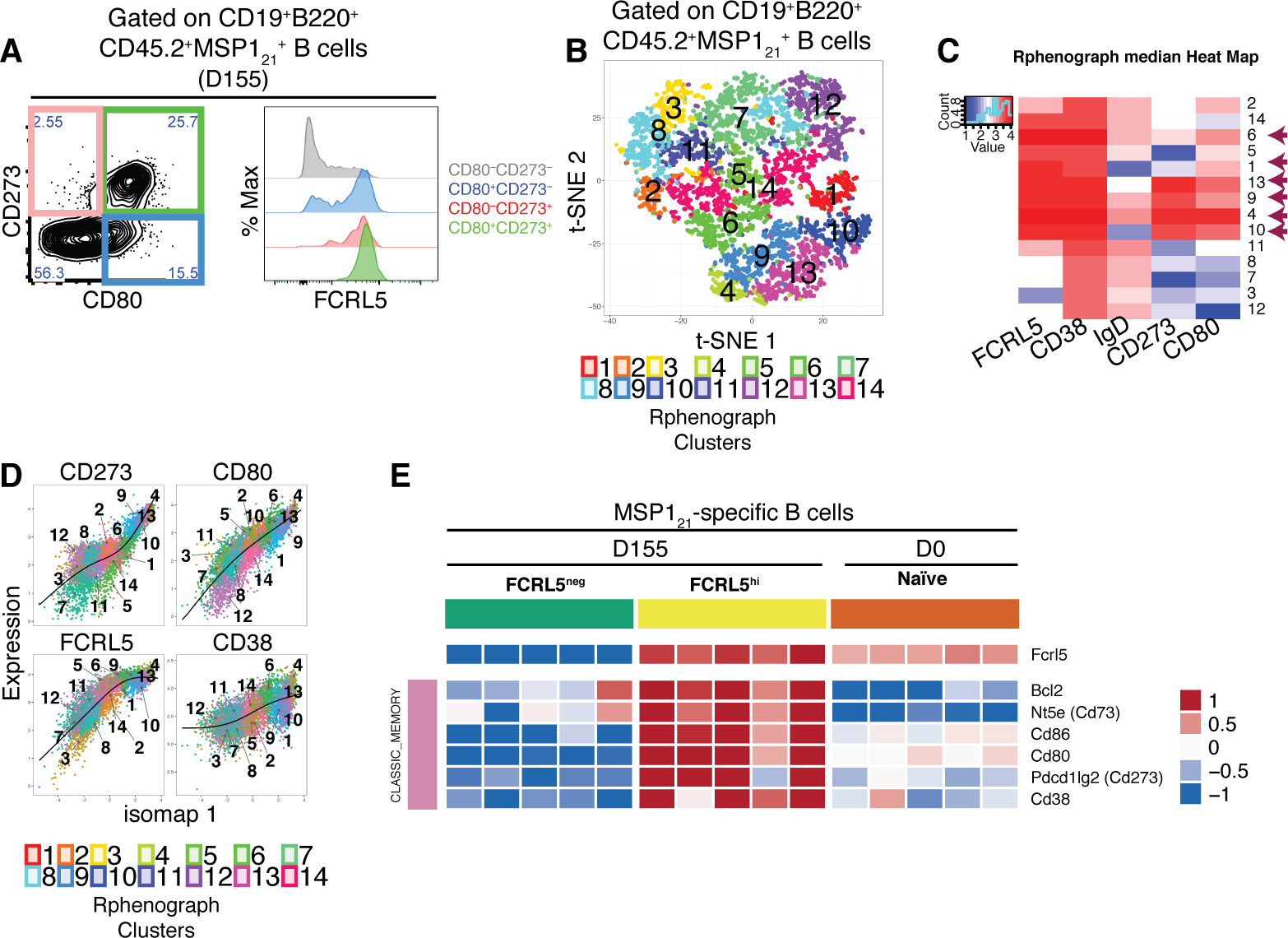
FCRL5^hi^ identifies MSP1_21_-specific B_mem_ after resolution of *P. chabaudi* infection. (A) Flow cytometry showing expression of FCRL5 (right) on different subsets of splenic MSP1_21_-specific B cells from NIMP23→*Rag2*^−/−^ chimeric mice defined based on CD273 and CD80 expression (left) at 155dpi. (B) t-SNE analysis of splenic MSP1_21_-specific B cells based on FCRL5, CD38, IgD, CD273 and CD80 expression measured by flow cytometry (n=5). Clusters identified by PhenoGraph are colored and numbered. (C) PhenoGraph heat map showing median expression of FCRL5, CD38, IgD, CD273 and CD80 on the different clusters of MSP1_21_-specific B cells. Arrows point at the different clusters displaying a memory B cell phenotype. (D) Expression profiles of FCRL5, CD38, CD273 and CD80 for the different PhenoGraph clusters visualized on the first component of ISOMAP. The regression line estimated using the generalized linear model (GLM) is added for each marker. Data representative of three independent experiments with 4-7 mice per group. (E) Heat map showing expression levels of different genes on splenic FCRL5^−^ and FCRL5^hi^ MSP1_21_-specific B cells sorted at 155dpi, and MSP1_21_-specific B cells sorted before infection (naive), determined by RNAseq analysis. Each column corresponds to data from an individual mouse (n=5 155dpi, n=5 0dpi).

### *Plasmodium*-specific AMB are a distinct short-lived activated B cell subset

After identifying and sorting MSP1_21_-specific AMB during chronic *P. chabaudi* infection, and MSP1_21_-specific B_mem_ after resolution of the infection, we then compared the transcriptome of these two B-cell subsets. Principal component analysis (PCA) demonstrated a strikingly distinct transcriptome of MSP1_21_-specific AMB from that of MSP1_21_-specific B_mem_, as well as all other MSP1_21_-specific B-cell subsets sorted in this study (Figure 7A). The MSP1_21_-specific AMB sorted at 35dpi formed a separated cluster at the extreme right of the PC1 axis of the PCA plot, which accounts for the majority of the variance (Figure 7A). All the other subsets [including MSP1_21_-specific CD11b^−^CD11c^−^ B-cells sorted from the same mice and at the same day post-infection as the MSP1_21_-specific AMB (i.e. 35dpi)] clustered on the left of the PC1 axis, and showed differences mostly along the PC2 axis of the PCA plot, which accounts for only 10% of the variance (Figure 7A). Interestingly, MSP1_21_-specific CD11b^−^CD11c^−^ and MSP1_21_-specific B_mem_ clustered on opposite sides of the MSP1_21_-specific naive B-cell subset along the PC2 axis (Figure 7A), which suggests that the MSP1_21_-specific B_mem_ more closely resemble MSP1_21_-specific naive B cells than MSP1_21_-specific CD11b^−^CD11c^−^ B cells sorted at 35dpi.

MSP1_21_-specific AMB sorted during chronic *P. chabaudi* infection, and MSP1_21_-specific B_mem_ sorted after resolution of the infection shared the expression of a series of mouse memory markers, namely *Cd80*, and also *Fcrl5*, *Nt5e* (CD73), and *Cd86* (Figure 7B). However, these two subsets showed differences in the expression pattern of anti-and pro-apoptotic genes (Figure 7C). While MSP1_21_-specific B_mem_ from after infection resolution showed the highest levels of expression of the anti-apoptotic *Bcl2* gene, MSP1_21_-specific AMB sorted during chronic *P. chabaudi* infection showed the lowest levels of expression of this hallmark anti-apoptotic gene (Figure 7C). In contrast to MSP1_21_-specific B_mem_, MSP1_21_-specific AMB expressed high levels of the pro-apoptotic genes *Bad*, *Bax*, *Fas* and *Fasl* (Figure 7C). Interestingly, MSP1_21_-specific AMB expressed very high levels of class-switched immunoglobulins, including *Igha*, *Ighg1*, *Ighg2b*, *Ighg2c* and *Ighg3* (Figure 7D). Finally, MSP1_21_-specific AMB highly expressed *Cd11b*, *Cd11c*, *Tbx21*, *Ifng* and *Pdcd1* (Figure 7E), all hallmarks of human AMB, as well as *Mki67*, indicative of active cell division, as previously shown in human AMB.

**Figure 7.**
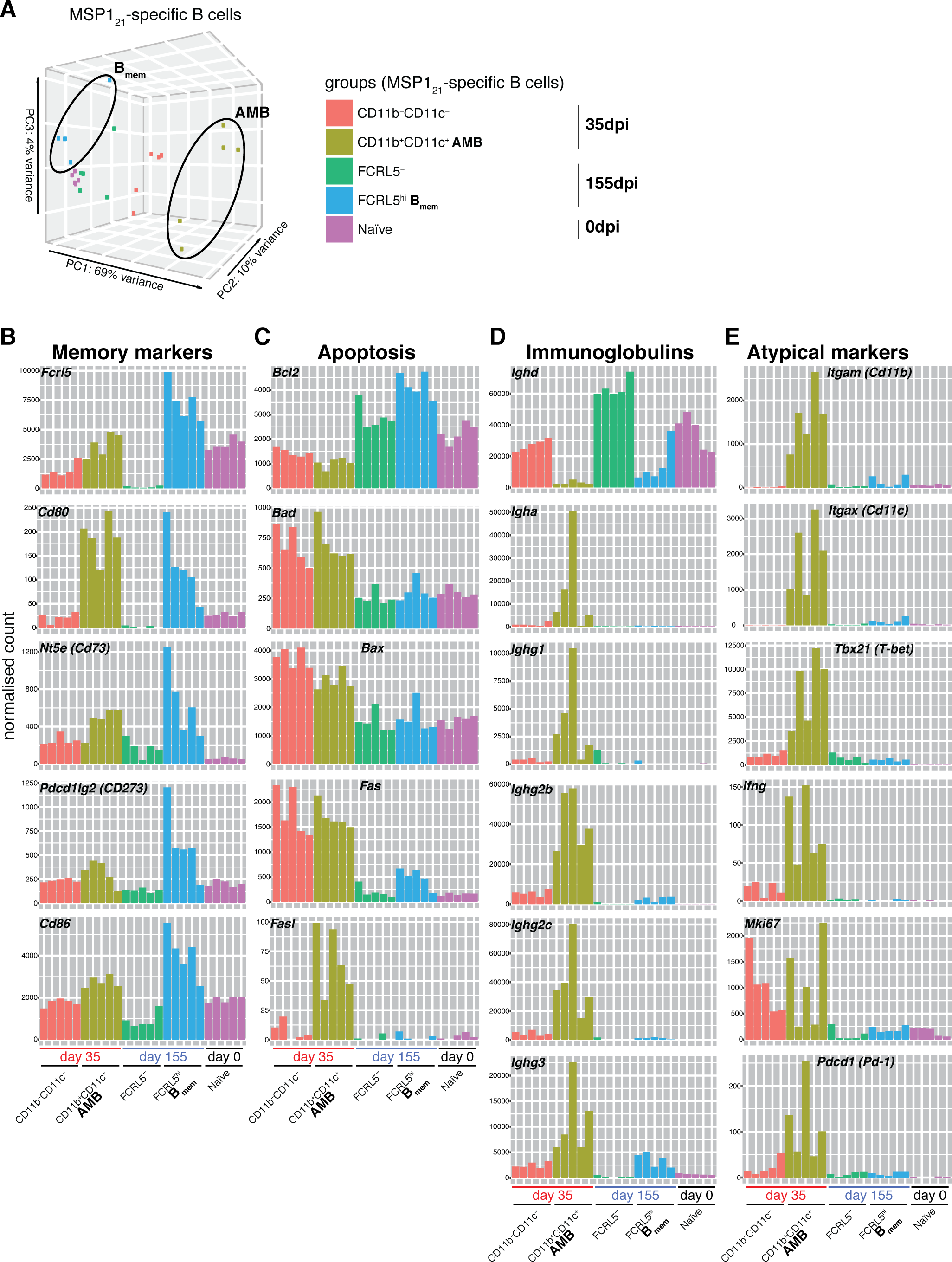
MSP1_21_-specific AMB are a distinct short-lived activated B cell subset. (A) Principal component analysis of RNAseq transcriptome data from splenic MSP121-specific AMB (CD11b^+^CD11c^+^, 35dpi), CD11b^−^CD11c^−^ B cells (35dpi), Bmem (FCRL5^hi^, 155dpi), FCRL5^−^ B cells (155dpi) and B cells from naïve mice (0dpi). The MSP1_21_-specific AMB and B_mem_ are contained inside ellipses. (B) (C) (D) (E) Normalized counts corresponding to selected genes representing memory B-cell markers, anti and pro-apoptotic genes, immunoglobulins and atypical memory B-cell markers, respectively, for all five groups described in (A). Each bar represents an individual mouse. Data generated with 5 mice per group.

These data demonstrate that AMB and B_mem_ share the expression of memory markers. However, they show striking differences in the expression of pro-and anti-apoptotic genes, immunoglobulins genes, and cell proliferation genes. The increased expression of *Mki67*, pro-apoptotic genes and class-switched immunoglobulins in AMB suggests that they resemble activated B cells. By contrast, B_mem_ express much less *Mki67* (similar to naïve cells) and present an anti-apoptotic gene expression pattern, consistent with being long-lived quiescent B cells.

## DISCUSSION

Similar to other chronic infections [e.g. HIV, HCV and *Mycobacterium tuberculosis* (Knox et al., 2017b; Portugal et al., 2017)], *Plasmodium* infection, the cause of malaria, leads to an increase in the frequency of AMB [originally termed tissue-like memory B cells (Moir et al., 2008)] in peripheral blood from *P. falciparum*-exposed subjects (Illingworth et al., 2013; Portugal et al., 2015; Sullivan et al., 2016; 2015; Weiss et al., 2011; 2010; 2009). However, mouse models to investigate *Plasmodium*-specific AMB are lacking. Here we have generated an IgH knock-in transgenic mouse strain to study the generation of *Plasmodium*-specific AMB in a *Plasmodium chabaudi* infection. We demonstrate the generation of *P. chabaudi*-specific AMB in response to blood-stage mosquito-transmitted chronic *P. chabaudi* infection, and show the short-lived nature of these cells. *P. chabaudi* infections in mice present some outstanding characteristics for the study of AMB; a chronic phase which allows investigation of the impact of constant immune activation driven by persistent subpatent parasitemia, followed by a clearance phase which allows the study of immune responses after the infection is naturally resolved. Thus, it is possible to study both exhaustion driven by chronic immune activation, and memory immune responses which remain after *P. chabaudi* elimination.

The *P. chabaudi*-specific AMB we detected during the chronic phase of infection showed strong similarities to human AMB described in chronic malaria. These included being class-switched, and expressing mouse homologues of hallmark human atypical memory B-cell genes such as *Itgax* (Cd11b), *Itgam* (Cd11c), *Cxcr3*, *Fcrl5*, *Tbx21* (T-bet), *Ifng*, *Cd80*, *Cd86*, *Aicda*, and a series on inhibitory receptors, including *Lair1* and *Pdcd1* (PD-1). Human AMB have been shown to share several characteristics with plasmablasts (Knox et al., 2017a; Muellenbeck et al., 2013; Obeng-Adjei et al., 2017; Sullivan et al., 2015). Interestingly, we also observed increased expression of several genes associated with plasmablasts in *P. chabaudi*-specific AMB, including *Cd138*, *Xbp1*, *Prdm1* (encoding Blimp-1) and *Mki67*, accompanied by downregulation of *Pax5* and *Bcl6*.

AMB resemble age-associated B cells (ABC) which accumulate with age as well as in autoimmunity, and were also proposed to be a subset of long-lived memory B cells (Naradikian et al., 2016a; Portugal et al., 2017; Rubtsov et al., 2017). Expression of T-bet, CD11c and CXCR3 are shared by AMB, tissue-like memory B cells, and ABC (Knox et al., 2017b; Naradikian et al., 2016a). Moreover, similar to ABC, expansion of human AMB associated with malaria is driven by IFNγ (Obeng-Adjei et al., 2017). IL-21, which is highly expressed by follicular helper T cells in response to *Plasmodium* infection (Carpio et al., 2015; Obeng-Adjei et al., 2015; Pérez-Mazliah et al., 2015), also directly promotes T-bet expression in B cells in the context of TLR engagement (Naradikian et al., 2016b). Taken together, these data strongly suggest that this is the same T-bet^+^ B-cell subset, which accumulates with time due to repetitive antigenic exposure. In agreement with previous data (Rubtsova et al., 2013), we show here that immunization with MSP1_21_ and R848, a TLR7/8 ligand, promotes a robust but short-lived CD11b^+^CD11c^+^ *P. chabaudi*-specific AMB response. T-bet^+^ atypical B cells are critical to eradicate various murine viral infections (Barnett et al., 2016; Rubtsova et al., 2013), and a recent study showed that yellow fever and vaccinia vaccinations of humans stimulated an acute T-bet^+^ B-cell response and suggested that these T-bet^+^ B-cell population may function as an early responder during acute viral infections (Knox et al., 2017a). Thus, T-bet^+^ B cells, even in the context of malaria, are likely to be a normal component of the immune compartment that becomes activated and expands, most probably in response to BCR, endosomal TLR, and IFNg or IL-21 stimulation. Moreover, a recent study shows a TLR9/IFNg-dependent activation of autoreactive T-bet^+^CD11c^+^ atypical B cells in response to *P. yoelii* 17XNL infection in mice (Rivera-Correa et al., 2017). However, whether these cells remained part of the long-lived memory B-cell pool after resolution of the infection was not explored.

Here, we show that *P. chabaudi*-specific AMB are short-lived activated B cells. These cells were absent after resolution of the infection, and immunization with purified antigen and TLR agonists resulted in a transient, yet robust, activation of *P. chabaudi*-specific AMB which lasted no more than 48h. Moreover, the *Igh*^NIMP23/+^ mouse model allowed us to obtain a deep insight of the transcriptome profile of MSP1_21_-specific AMB during natural infection, and compare it side-by-side with the transcriptome of MSP1_21_-specific naïve B cells. This allowed us to demonstrate their heavily pro-apoptotic and activated transcription profile, further explaining their short-lived nature. *P. chabaudi*-specific AMB showed very low expression of *Bcl2*, and high levels of expression of several pro-apoptotic genes including *Bad*, *Bax*, *Fas* and *Fasl*. In addition, these cells expressed very high levels of class-switched immunoglobulins and genes associated with DNA replication and proliferation.

The association of *P. chabaudi*-specific AMB with ongoing infection explains several observations in human studies: reduction of HIV plasma viremia by ART resulted in a significant reduction of HIV-specific AMB without altering the frequency of HIV-specific B_mem_ (de Bree et al., 2017; Kardava et al., 2014); individuals living in high malaria endemicity present higher frequencies of AMB than individuals living in areas with moderate transmission (Illingworth et al., 2013; Sullivan et al., 2015); repetitive *Plasmodium* episodes result in higher frequencies of AMB (Obeng-Adjei et al., 2017); the percentage of AMB is larger in children with persistent asymptomatic *Plasmodium falciparum* parasitemia as compared with parasite-free children (Weiss et al., 2009); previously exposed subjects significantly reduce the frequency of AMB following a year of continuous absence of exposure to *Plasmodium falciparum* infection (Ayieko et al., 2013). These observations all support the view that constant immune activation rather than impaired memory function leads to the accumulation of AMB in malaria.

After resolution of infection, *P. chabaudi*-specific AMB did not persist, but instead subsets of *P. chabaudi*-specific B_mem_ were readily detected. These cells expressed different combinations of previously described mouse B-cell memory markers [i.e. CD80, CD273 and CD73 (Anderson et al., 2007; Tomayko et al., 2010; Zuccarino-Catania et al., 2014)]. *P. chabaudi*-specific B_mem_ included both class-switched and non-class-switched cells, which show different responses to a secondary challenge infection (Krishnamurty et al., 2016). Independently of the combination of previously described memory markers expressed on *P. chabaudi*-specific B_mem_, all of these cells displayed very high expression of FCRL5. Previous data showed most prominent expression of FCRL5 in marginal zone B cells, while much less evident in the newly-formed and follicular splenic B cell subpopulations (Davis, 2004; Won et al., 2006). In agreement with this, we observed expression of FCRL5 on a subset of splenic *P. chabaudi*-specific B cells obtained from naïve mice. However, the level of FCRL5 expression on *P. chabaudi*-specific B_mem_ detected after resolution of the infection was noticeably higher than that of naïve B cells both at the protein and mRNA levels. In humans, FCRL4 has been shown to be preferentially expressed on memory B cells (Ehrhardt et al., 2005; 2003). Similar to expression on human B cells, we show that members of the FCRL family are expressed on mouse B_mem_. In contrast to MSP1_21_-specific AMB, these CD11b^−^CD11c^−^FCRL5^hi^ MSP1_21_-specific B_mem_ showed high expression of the hallmark memory and anti-apoptotic gene *Bcl2* (Bhattacharya et al., 2007). Thus, after resolution of the infection, high expression of FCRL5 acted as a universal B_mem_ marker. We further confirmed the high expression of FCRL5 on MSP1_21_-specific B_mem_ generated by immunization with model antigen. However, as *P. chabaudi*-specific AMB also expressed CD80 and high levels of FCRL5, we believe these proteins are general markers of antigen-experienced B cells. Nonetheless, due to its complex dual ITIM/ITAM signaling capacity (Zhu et al., 2013), it is tempting to speculate that FCRL5 might serve as an important signal in the differentiation/maintenance of B_mem_.

Tracking the fate of the different MSP1_21_-specific B-cell subsets identified in this work will allow detailing the interplay between them. A model can be proposed in which antigen-specific AMB serve as an intermediate stage of differentiation between naïve and B_mem_. Alternatively, antigen-specific AMB might represent early plasmablasts. These two scenarios are not necessarily antagonist, and might even occur in parallel.

Our data suggest that the expansion of AMB in malaria is not a consequence of B-cell exhaustion, but rather a physiologic stage of B-cell activation, and that these cells are sustained in high frequencies by ongoing chronic infections. Thus, *Plasmodium*-specific AMB are neither “memory”, nor “atypical”. Importantly, our data demonstrate that robust expansion of *Plasmodium*-specific AMB does not hinder clearance of the infection, activation of germinal centers, or generation of *Plasmodium*-specific long-lived quiescent B_mem_ upon resolution of the infection.

## MATERIALS AND METHODS

### Mice

5-12week-old female mice were used for experiments. C57BL/6J, C57BL/6.SJL-*Ptprc*^*a*^ (CD45.1 congenic), *Rag2*^−/−^. C57BL/6.SJL-*Ptprc*^*a*^ (CD45.1 congenic) and BALB/c mouse strains were bred in the specific pathogen-free facilities of the MRC National Institute for Medical Research and The Francis Crick Institute, and were housed conventionally with sterile bedding, food and irradiated water. Room temperature was 22°C with a 12h light/dark cycle; food and water were provided ad libitum. The study was carried out in accordance with the UK Animals (Scientific Procedures) Act 1986 (Home Office license 80/2538 and 70/8326), was approved by the MRC National Institute for Medical Research Ethical Committee and was approved by The Francis Crick Institute Ethical Committee.

To produce MSP1_21_specific B cell knock-in mice capable of undergoing class switch recombination on the C57BL/6J genetic background, the VDJ_H_^NIMP23^ anti-MSP1_21_ variable region coding exon containing the Leader-V segment intron from gDNA of the NIMP23 hybridoma (Boyle et al., 1982) was inserted by homologous recombination into the 5’ end of the endogenous IgH locus (Taki et al., 1993) Figure S1A-C). The VDJ_H_^NIMP23^ anti-MSP1 21 variable region coding exon was inserted into a previously described IgH targeting construct, replacing the anti-HEL heavy chain variable region coding exon that was already in it (Phan et al., 2003) (Figure S1D). The final targeting construct included a loxP-flanked neomycin resistance cassette in reverse transcriptional orientation to the *Igh* locus, located immediately 5’ to the rearranged VDJ_H_^NIMP23^ variable region and its associated promoter (Figure S1D).

Electroporation of C57BL/6N-derived PRX embryonic stem cells with the targeting construct and selection of homologous recombinant clones was performed using standard techniques by PolyGene AG (Switzerland). One targeted ES clone was used for production of chimeric mice using standard techniques at the Biological Research Facilities of the MRC National Institute for Medical Research, London, UK. Male chimeric mice were crossed to C57BL/6J females and progeny carrying the *Igh*^NIMP23neo^ allele were crossed to PC3Cre mice (O’Gorman et al., 1997) to delete the neo^r^ gene in the germline and generate mice carrying the *Igh*^NIMP23^ allele (Figure S1D). The *Igh*^NIMP23/+^ strain was maintained by backcrossing for at least 10 generations to C57BL/6J mice.

### Mixed bone marrow chimeras

Femurs and tibias were excised from female mice and cleaned of flesh using forceps and scalpel, and BM was obtained by flushing out with IMDM supplemented with 2 mM L-glutamine, 0.5 mM sodium pyruvate, 100 U penicillin, 100 mg streptomycin, 6 mM Hepes buffer, and 50mM 2-ME (Gibco, Invitrogen), using a syringe with a needle. Thereafter, single BM cell suspensions were obtained by mashing through a 70μm filter mesh, further sieved through 40μm filter mesh and washed once. Live cells were resuspended in sodium chloride solution 0.9% (Sigma) at 4×10^6^ cells/200μl. *Rag2*^−/−^.C57BL/6.SJL-*Ptprc*^*a*^ mice were sub-lethally irradiated (5Gy) using a [137Cs] source and reconstituted less than 24hr after irradiation by i.v. injections of a 10% *Igh*^NIMP23/+^:90% C57BL/6.SJL-*Ptprc*^*a*^ combination of donor BM cells. Recipient mice were maintained on acidified drinking water and analyzed for reconstitution after 6-8 weeks.

### *Plasmodium chabaudi* infection

*Plasmodium chabaudi chabaudi AS* was transmitted by *Anopheles stephensi* mosquitoes, strain SD500, as described elsewhere (Spence et al., 2012). Briefly, C57BL/6J mice were injected i.p. with 10^5^ *P. chabaudi*-infected red blood cells and used to feed mosquitos two weeks after the injection. Two weeks after mosquito feeding/infection, each experimental mouse was exposed to 20 infected mosquitos for 30 min. Blood parasitemia in infected experimental mice was routinely monitored by thin blood smears.

### Immunizations

Mice were immunized i.p. with a combination of 100μg of MSP1_21_ (Quin and Langhorne, 2001) and 50μl of Titermax Gold emulsion (Sigma), or a combination of 50 μg of MSP1_21_ and 50 μg of R848 (Invivogen).

### Flow cytometry and cell sorting

Spleens, lymph nodes and bone marrows were dissected and single cell suspensions were obtained by mashing the organs through a 70μm filter mesh in HBSS, 6mM Hepes buffer (Gibco, Invitrogen). After removal of red blood cells from spleens and bone marrows by treatment with lysing buffer (Sigma), the remaining cells were resuspended in complete Iscove’s Modified Dulbecco’s Medium [IMDM supple-mented with 10% FBS Serum Gold (PAA Laboratories, GE Healthcare), 2 mM L-glutamine, 0.5 mM sodium pyruvate, 100U penicillin, 100 mg streptomycin, 6 mM Hepes buffer, and 50 mM 2-ME (all from Gibco, Invitrogen)] and viable cells were counted using trypan blue (Sigma) exclusion and a hemocytometer. Cells were then resuspended in PBS and incubated with APC-or PE-labelled MSP1_21_ fluorescent probes and/or different combinations of fluorochrome-conjugated antibodies (Table S3), and either acquired after two washes with PBS, or fixed with 2% paraformaldehyde and stored in staining buffer at 4°C until acquisition.

The APC and PE MSP1_21_ fluorescent probes were produced as previously described (Krishnamurty et al., 2016; Martinez et al., 2012). Briefly, purified MSP1_21_ (Quin and Langhorne, 2001) was biotinylated using an EZ-link Sulfo-NHS-LC-Biotinylation kit (Thermo Fisher Scientic) using a 1:1 ratio of biotin to protein, and loaded onto Streptavidin-APC conjugated or Phycolink Streptavidin-R-PE conjugated (ProZyme) in a 6:1 ratio of MSP1_21_:Streptavidin-fluorochrome.

Cell sorting was performed on a MoFlo XDP (Beckman Coulter) or a BD FACSAria Fusion (BD Biosciences) and the target cell populations were directly dispensed into TRIreagent (Ambion) and stored at −80°C until RNA isolation. Purity checks were routinely performed for all assays by sorting aliquots of cells into PBS containing 2% FCS and reacquiring them on the cell sorter.

Dead cells were routinely excluded from the analysis by staining with LIVE/DEAD Fixable Aqua or Blue stain (Invitrogen). Singlets were selected based on FCS-A vs FCS-H and further based on SSC-A vs SSC-H. “Fluorescence minus one” (FMO) controls were routinely used to verify correct compensation and to set the thresholds for positive/negative events. Analysis was performed with FlowJo software version 9.6 or higher (Tree Star).

PhenoGraph and *t*-distributed stochastic neighbor embedding (t-SNE) were combined to analyze multiparameter flow cytometry data using the Cytofkit package (Chen et al., 2016). t-SNE renders high-dimensional single-cell data based on similarities into only two dimensions, and thus helps visualize multiparameter data (van der Maaten, 2008). PhenoGraph (Levine et al., 2015) allows partitioning of highdimensional single-cell data into phenotypically coherent subpopulations (i.e. clusters). The relatedness of the cell clusters identified by PhenoGraph was inferred using Isomap (Cytofkit package), in which related clusters/subsets can be visualised close to each other.

### RNA isolation, sequencing and data analysis

Total RNA from 1-5×10^4^ cells sorted into TRIreagent (Ambion) was isolated using the Ribopure kit (Ambion). Concentration of purified RNA was determined by Qubit fluorometric quantitation using the HS assay kit (ThermoFisher Scientific), and the quality analyzed with a 2100 Bioanalyzer (Agilent). Samples with a RIN score above 8.50 were used for the next steps. cDNA was generated from total RNA with the SMART-Seq v4 Ultra Low Input RNA Kit (Takara Bio USA). Next-generation sequencing libraries were produced with the Ovation Ultralow System V2 (Nugen), and run as PE100 on a HiSeq 4000 sequencer (Illumina). GEO accession: GSE115155.

For bioinformatics analysis, paired-end sequence reads were adapter and quality trimmed using cutadapt v1.9.1 (Martin, 2011) with the following non-default settings: “-a AGATCGGAAGAGC-A AGATCGGAAGAGC --minimum-length 30-q 20,20”. Gene-level abundance estimates were generated from the trimmed reads using RSEM v1.2.31 (B. Li and Dewey, 2011) running STAR v2.5.1b (Dobin et al., 2013) with default settings, aligned against the *Mus musculus* Ensembl release 89 transcriptome (mm10). All further analysis was conducted using the DESeq2 (Love et al., 2014) package from Bioconductor v3.5 run in R v3.4.0. The expected counts were imported and rounded to integers to generate a counts matrix. Differential expression between phenotype groups was assessed using the DESeq function with default settings. In the case of comparisons of different MSP1_21_-specific B cell subsets obtained from the same experimental mouse, an additional mouse factor was added to the design formula to accommodate the paired nature of the data. Significance was thresholded using an FDR≤0.01. PCA analysis was conducted using DESeq’s plot PCA function with the regularized log (rlog) transformed count data. Heat maps were generated using the regularized log (rlog) transformed count data, scaled per gene using a z-score. Significant genes with a normalized read count consistently ≥100 in all samples from at least one of the replicate groups being compared were ranked according to the fold changes and use to generate the top 50 most up and down-regulated gene heat maps. Significant selected marker genes were used to produce separate heat maps split by functional annotation. The GSEA pre-ranked function from the Broad’s Gene Set Enrichment Analysis (GSEA) (Subramanian et al., 2005) suite was used to assess significant enrichment of MSigDB’s C2 Reactome gene sets associated with differential expression between cell types. The function was run using a list of genes ranked for differential expression using DESeq2’s Wald test statistic with default settings except for:

collapse dataset to gene = false
enrichment statistic = classic

### Statistical analysis

Statistical analysis was performed using Mann Whitney U test, Kruskal-Wallis test followed by Dunn’s multiple comparisons test, or Two-Way ANOVA followed by Dunnett’s multiple comparisons test on Prism software version 6 (GraphPad). P<0.05 was accepted as a statistically significant difference.

## AUTHOR CONTRIBUTIONS

Conceived the study: DPM VT JL. Designed experiments: DPM PG JL ES VT. Wrote the manuscript: DPM VT JL. Managed transgenic mouse colonies: DPM AP SM. Performed experiments: DPM PG CH SM IT. Analyzed data: DPM PG. Supervised the study: VT JL.

## ACKNOWLEDGEMENTS

We are grateful to Graham Preece, Philip Hobson and the Flow Cytometry Facility, Jackie Holland and the Biological Research Facility, Richard Mitter and the Bioinformatics Facility, and the Advanced Sequencing Facility at the Francis Crick Institute and the former MRC National Institute for Medical Research, London, UK. We thank Marion Pepper, University of Washington, for her support with the preparation of the MSP1_21_ tetramers, and Dinis Calado and Matthew Lewis, Francis Crick Institute, for their critical reading of the manuscript.

This work was supported by the Francis Crick Institute which receives its core funding from the UK Medical Research Council (FC001101), Cancer Research UK (FC001101) and the Wellcome Trust (FC001101); by the Wellcome Trust (grant reference WT101777MA). Irene Tumwine is the recipient of a Francis Crick PhD studentship.

All authors, no conflict of interest.

---

**Figure S1.**
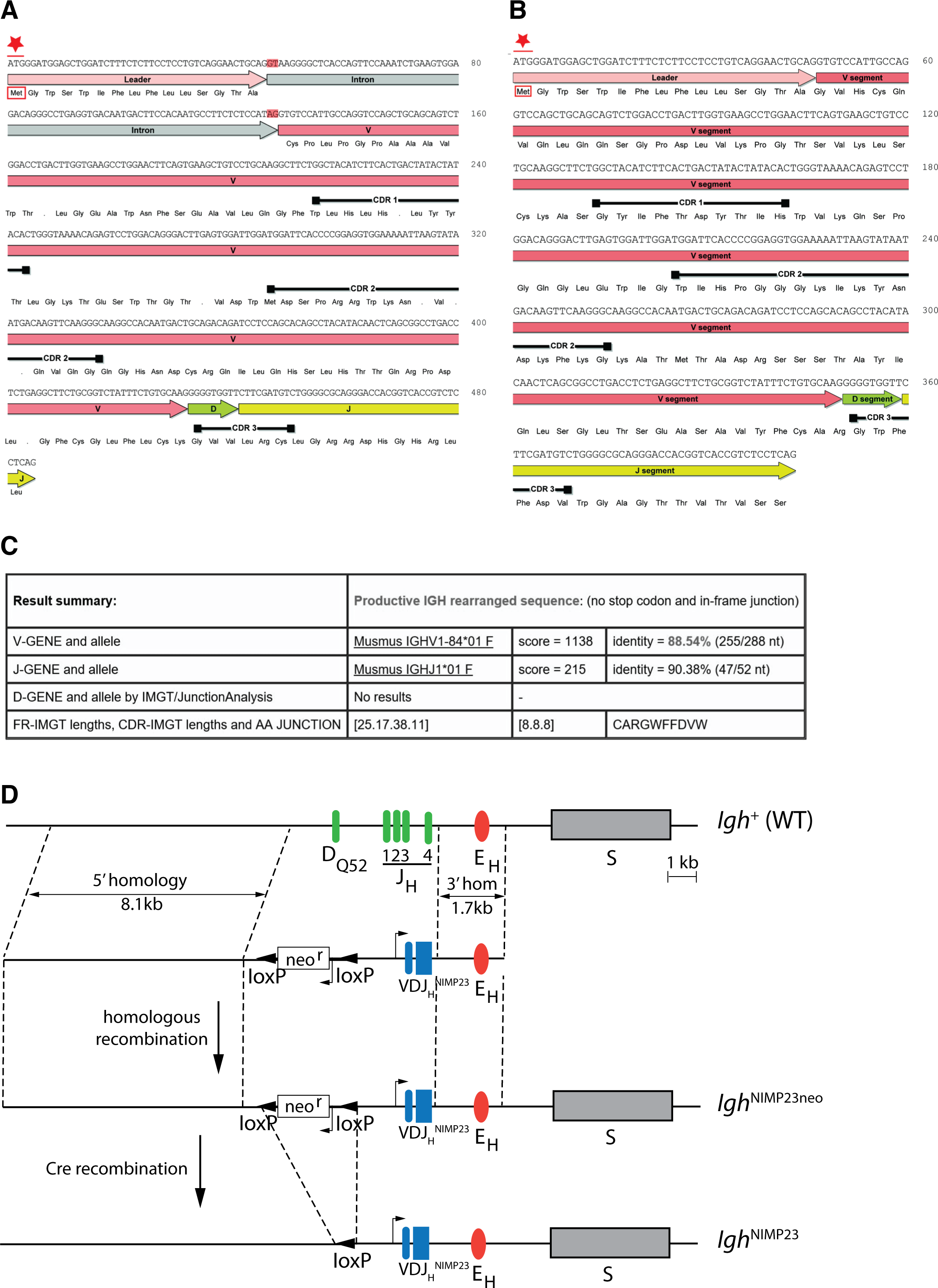
Generation of *Igh*^NIMP23/+^ knock-in mice. Annotated DNA and corresponding amino acid sequence of VDJ_H_^NIMP23^ obtained from (A) gDNA including the Leader-V intron and (B) cDNA, of the NIMP23 hybridoma: start Methionine (Met) indicated by red star, intron splice donor and acceptor sites highlighted in red, dots indicate STOP codons, black bars indicate predicted complementarity determining regions according to the Kabat database (Johnson and Wu, 2001) (C) IMGT/V-Quest mouse Ig database (Lefranc et al., 1999) comparative analysis result summary of the gDNA derived VDJ_H_^NIMP23^ sequence reveals the identity of the closest matching endogenous V and J genes (D) Schematic representation of (*Igh*^+^) Endogenous IgH locus showing the 4 JH segments, the DQ52 element, the Igh intronic enhancer (E_H_), the switch region for the constant μ gene (S), and (below) targeting construct indicating 5’ and 3’ homology arms and the inverted loxP-neo^r^-loxP cassette and rearranged VDJ_H_^NIMP23^ variable heavy chain region gene of the NIMP23 hybridoma, replacing DQ52 and all four JH segments of the endogenous IgH gene. *Igh*^NIMP23neo^: Targeted *Igh* locus after homologous recombination. *Igh*^NIMP23^: Final allele after Cre-mediated removal of neo^r^.

**Figure S2.**
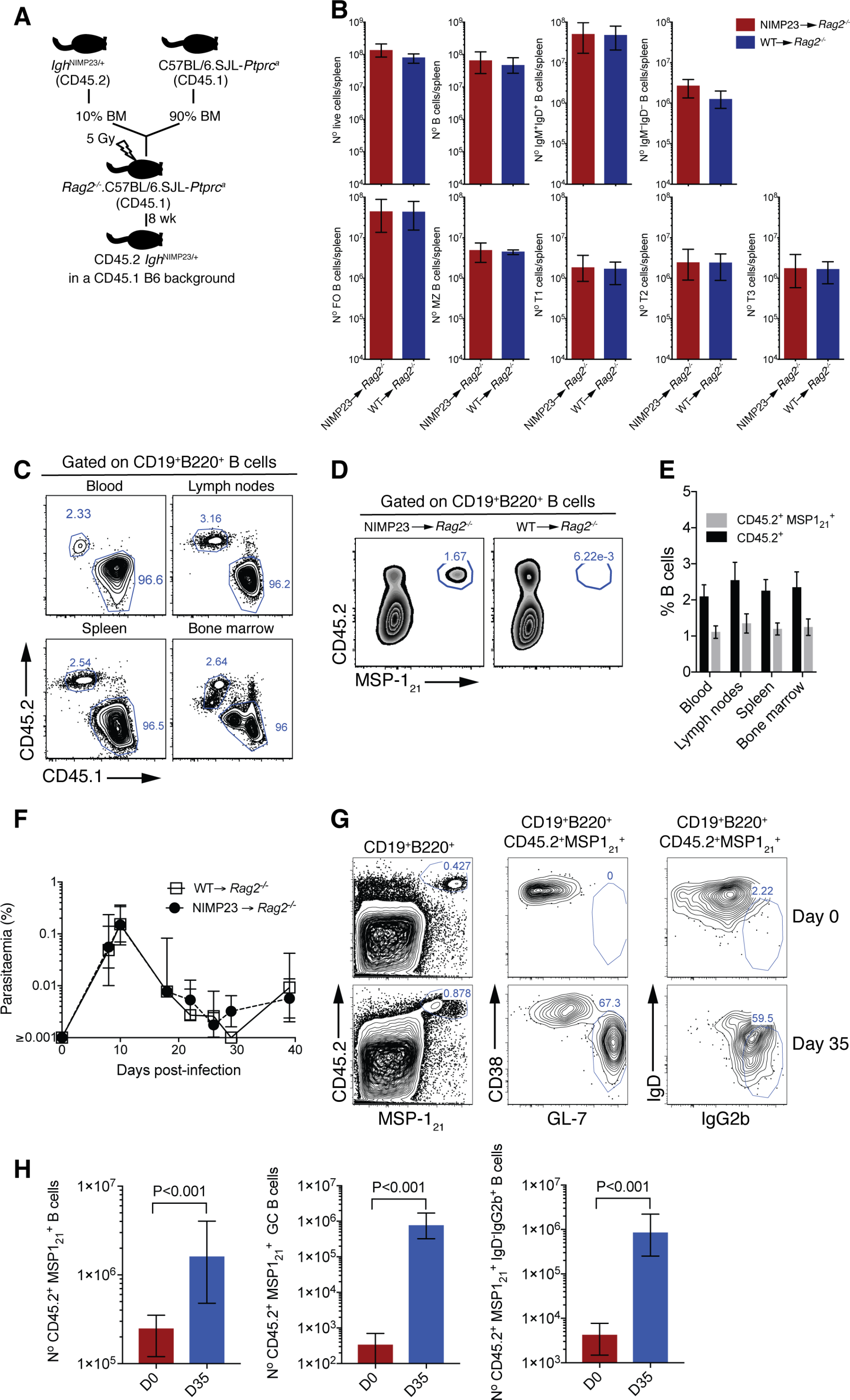
Generation of mixed bone marrow chimera model with reduced precursor frequency of *Igh*^NIMP23/+^ B cells to study MSP1_21_-specific B cell responses during *P. chabaudi* infection. (A) Experimental strategy to generate mixed bone marrow chimeric mice. (B) Numbers of different splenic B-cell populations defined by flow cytometry in *Rag2*^−/−^.C57BL/6.SJL-*Ptprc*^*a*^ mice reconstituted with a mixture of *Igh*^NIMP23/+^ and C57BL/6.SJL-*Ptprc*^*a*^ bone marrow in a 10:90 ratio (NIMP23→*Rag2*^−/−^), and control mice reconstituted with C57BL/6.SJL-*Ptprc*^*a*^ bone marrow (WT→*Rag2*^−/−^). Mann Whitney U test. Error bars are SEM. Data are representative of two independent experiments with 5 mice per group. (C) Flow cytometry of B cells obtained from different tissues of NIMP23→*Rag2*^−/−^ chimeric mice. Gates show frequencies of CD45.1^+^CD45.2^−^ and CD45.1^−^CD45.2^+^ (D) Flow cytometry of B cells obtained from spleen of NIMP23→*Rag2*^−/−^ and WT→*Rag2*^−/−^ control chimeric mice. Gates show frequencies of MSP1_21_-specific B cells as determined by CD45.2 vs MSP1_21_ staining. (E) Frequencies of CD45.1^−^CD45.2^+^ (black) and CD45.2^+^MSP1_21_^+^ (grey) B cells as gated in C and D, obtained from different organs of NIMP23→*Rag2*^−/−^ chimeric mice. (F) Blood-stage *P. chabaudi* parasitemia following mosquito transmission in NIMP23→*Rag2*^−/−^ and WT→*Rag2*^−/−^ control chimeric mice. (G) Flow cytometry data showing frequencies of MSP1_21_-specific GC (CD38^lo^GL-7^hi^) and class-switched (IgD^−^ IgG2b^hi^) B cells in the spleen of NIMP23→*Rag2*^−/−^ chimeric mice before infection (day 0) and at day 35 post-mosquito transmitted *P. chabaudi* infection. (H) Numbers of MSP1_21_-specific B cells, GC and class-switched B cells in the spleen of NIMP23→*Rag2*^−/−^ chimeric mice as gated in B and E. Mann Whitney U test. Error bars are SEM. Data representative of two independent experiments with 3-7 mice per group.

**Figure S3.**
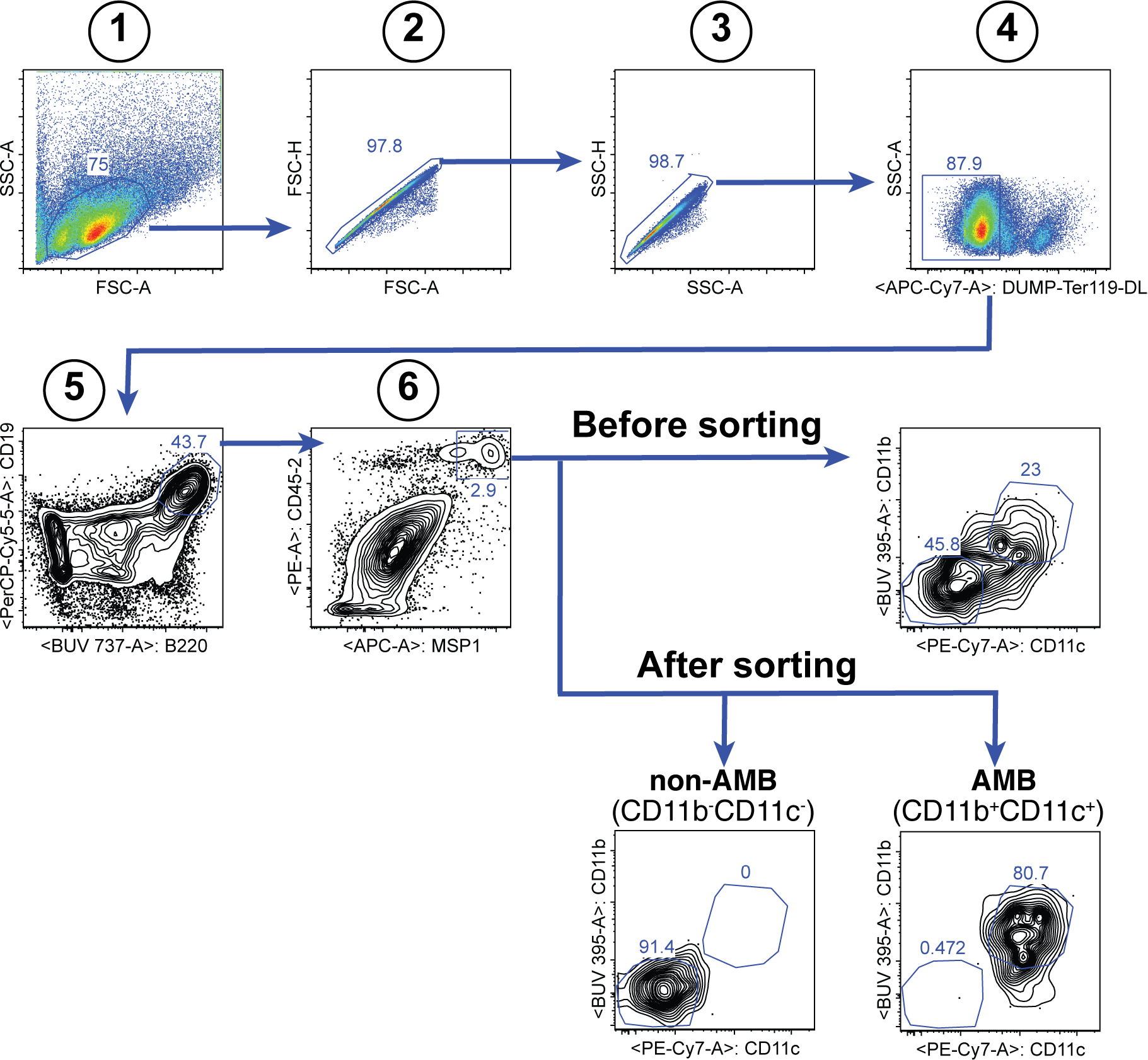
Gating strategy for the sorting of splenic MSP121-specific CD11b^+^CD11c^+^ AMB and CD11b^−^CD11c^−^ B cells at 35dpi.

**Figure S4.**
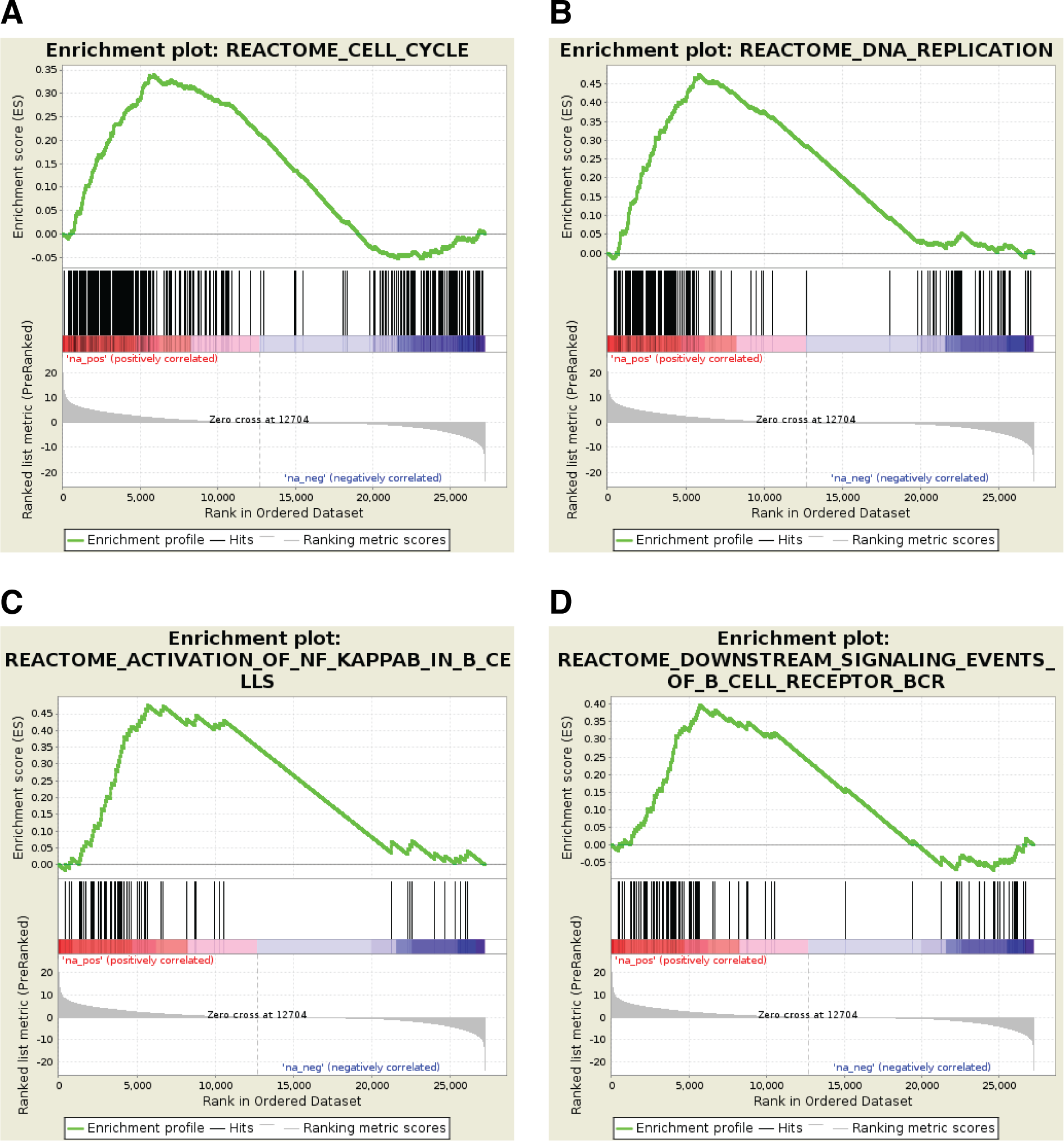
Profile of the Running *ES* Score and positions of GeneSet members on the rank ordered list for selected Reactome pathway gene sets.

**Figure S5.**
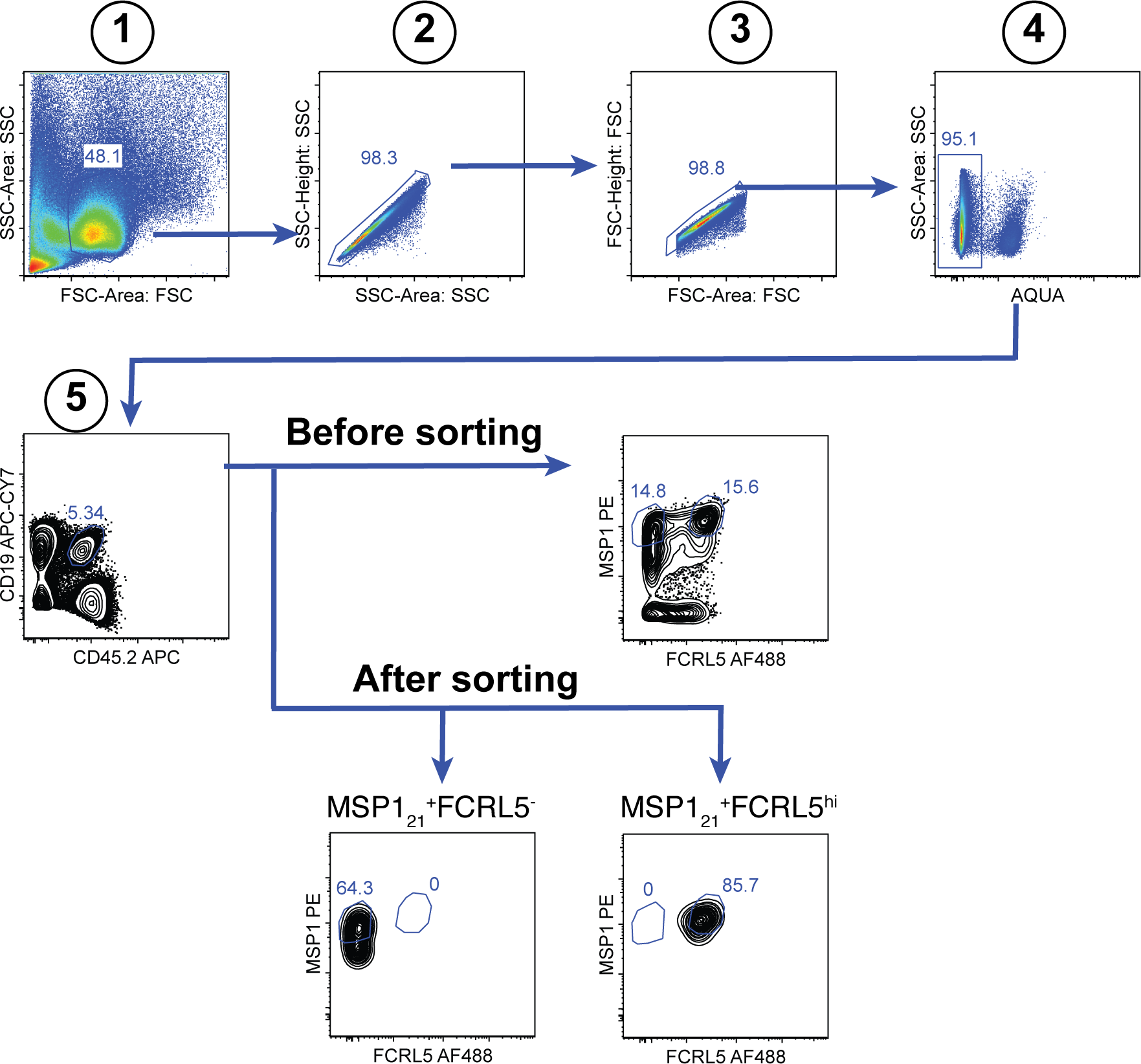
Gating strategy for the sorting of splenic MSP1_21_-specific FCRL5^hi^ B_mem_ and FCRL5^−^ B cells at 155dpi.

**Figure S6.**
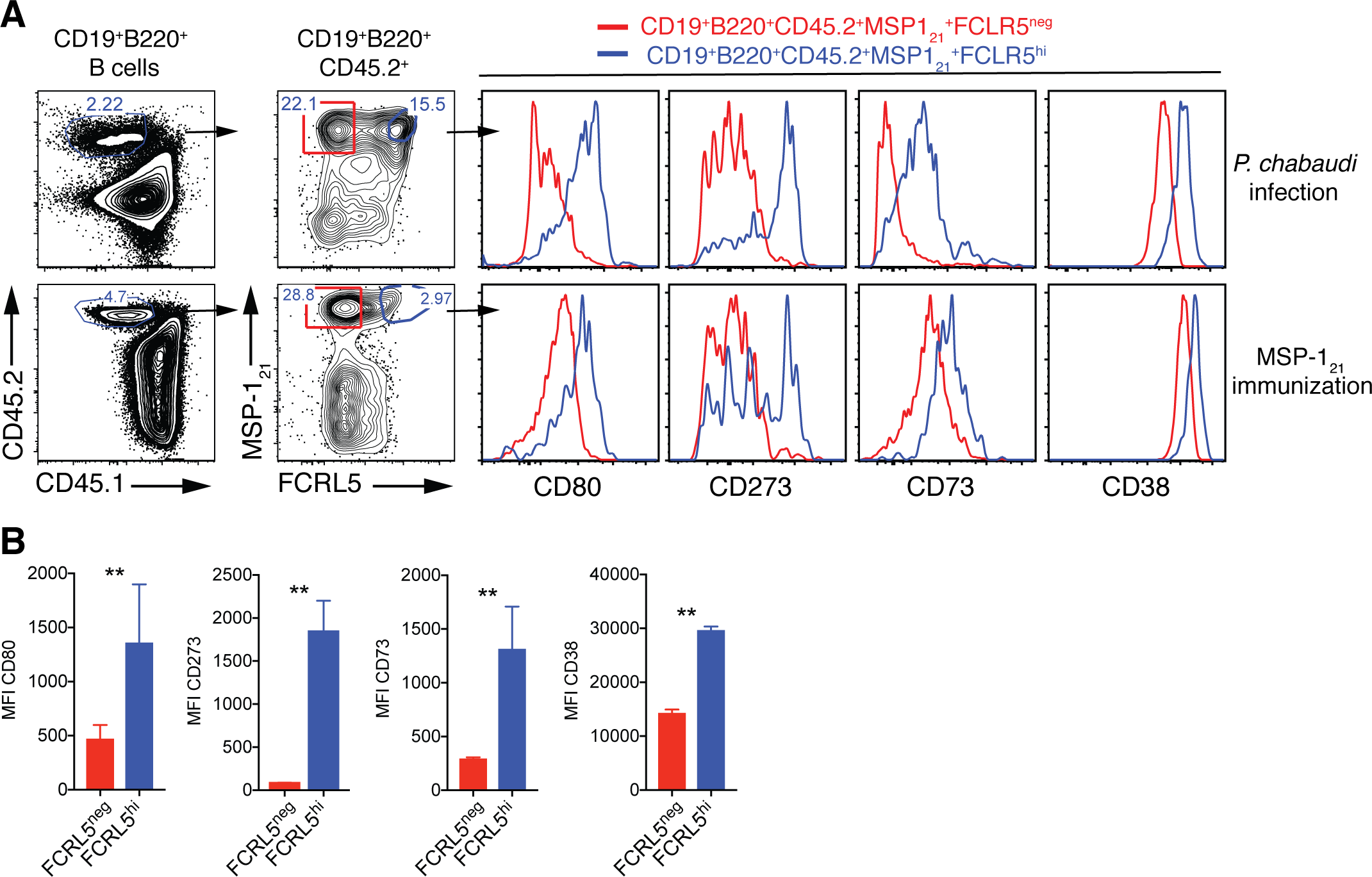
High expression of FCRL5 identifies B_mem_. (A) Flow cytometry gating strategy to identify splenic MSP1_21_-specific FCRL5^neg^ (red) and FCRL5^hi^ (blue) B cells, and additional surface markers expressed on these cells, in NIMP23→*Rag2*^−/−^ mixed bone marrow chimeras infected with *P. chabaudi* (155dpi, top row) and immunized with recombinant MSP1_21_ in Titermax Gold emulsion (day 29 post-immunization, bottom row). (B) Cumulative data showing geometric MFIfor CD80, CD273, CD73, and CD38 memory markers on splenic MSP1_21_-specific FCRL5^neg^ and FCRL5^hi^ B cells obtained from immunized mice. Mann Whitney U test. **, P<0.01. Error bars are SEM. Data representative of two independent experiments with 3-5 mice per group.

**Table S1.** Mouse homologues to human genes previously described to be either up (↑) or down (↓) regulated in human AMB.

| Gene | Common name | Reference | Functional group | AMB | Mouse homologue |
| --- | --- | --- | --- | --- | --- |
| MS4A1 | CD20 | MOIR 2008, SULLIVAN 2015 | **SURFACE PHENOTYPE** | ↓ | Ms4a1 |
| CR2 | CD21 | MOIR 2008, SULLIVAN 2015, PORTUGAL 2015 | **SURFACE PHENOTYPE** | ↓ | Cr2 |
| CD24A | CD24 | MULLENBECK 2013, SULLIVAN 2015 | **SURFACE PHENOTYPE** | ↓ | Cd24a |
| CD40 | CD40 | LI 2016, PORTUGAL 2015 | **SURFACE PHENOTYPE** | ↓ | Cd40 |
| ICOSL | ICOSL | LI 2016, | **ACTIVATION** | ↓ | Icosl |
| CD80 | CD80 | MOIR 2008, | **ACTIVATION** | ↑ | Cd80 |
| CD86 | CD86 | MOIR 2008, PORTUGAL 2015 | **ACTIVATION** | ↑ | Cd86 |
| FASL | CD95 | MOIR 2008, CHARLES 2011, | **ACTIVATION** | ↑ | Fasl |
| CD38 | CD38 | MOIR 2008, | **ACTIVATION** | ↑ | Cd38 |
| MKI67 | Ki-67 | MOIR 2008, MUELLENBECK 2013 | **ACTIVATION** | ↑ | Mki67 |
| FCGR2B | CD32 | SULLIVAN 2015, KARDAVA 2011 | **INHIBITORY** | ↑ | Fcgr2b |
| CD22 | CD22 | MOIR 2008, | **INHIBITORY** | ↑ | Cd22 |
| CD84 | CD84 | KARDAVA 2014, | **INHIBITORY** | ↑ | Cd84 |
| LILRB1 | CD85J | MOIR 2008, SULLIVAN 2015, PORTUGAL 2015 | **INHIBITORY** | ↑ | Lilrb1 |
| LILRB4A | CD85K | MOIR 2008, SULLIVAN 2015, PORTUGAL 2015 | **INHIBITORY** | ↑ | Lilrb4a |
| CD72 | CD72 | MOIR 2008, | **INHIBITORY** | ↑ | Cd72 |
| LAIR1 | LAIR-1 | MOIR 2008, | **INHIBITORY** | ↑ | Lair1 |
| PDCD1 | PD-1 | KARDAVA 2011, SULLIVAN 2015, PORTUGAL 2015 | **INHIBITORY** | ↑ | Pdcd1 |
| SIGLEC5 | SIGLEC5 | KARDAVA 2011, SULLIVAN 2015, PORTUGAL 2015 | **INHIBITORY** | ↑ | Siglece |
| SIGLEC6 | SIGLEC6 | KARDAVA 2011, SULLIVAN 2015, PORTUGAL 2015 | **INHIBITORY** | ↑ | Siglecf |
| LILRB3 | LILRB-3 | PORTUGAL 2015, | **INHIBITORY** | ↑ | Pirb |
| CXCR3 | CXCR3 | MOIR 2008, CHARLES 2011, SULLIVAN 2015 | **TRAFFICKING** | ↑ | Cxcr3 |
| CXCR5 | CXCR5 | MOIR 2008, SULLIVAN 2015 | **TRAFFICKING** | ↓ | Cxcr5 |
| CCR6 | CCR6 | MOIR 2008, SULLIVAN 2015 | **TRAFFICKING** | ↓ | Ccr6 |
| CCR7 | CCR7 | MOIR 2008, PORTUGAL 2015 | **TRAFFICKING** | ↓ | Ccr7 |
| ITGAX | CD11C | MOIR 2008, CHARLES 2011, | **TRAFFICKING** | ↑ | Itgax |
| ITGAM | CD11B | MOIR 2008, | **TRAFFICKING** | ↑ | Itgam |
| SELL | CD62L | MOIR 2008, SULLIVAN 2015 | **TRAFFICKING** | ↓ | Sell |
| FCRL3 | FCRL3 | MOIR 2008, SULLIVAN 2015, PORTUGAL 2015 | **FCRLs** | ↑ | Fcrl5 |
| TBX21 | TBET | RUSSELL KNODE 2017, MOIR 2008, SULLIVAN 2015, KNOX 2017 | **TRANSCRIPTION FACTORS** | ↑ | Tbx21 |
| IFNG | IFNγ | RUSSELL KNODE 2017, MOIR 2008, SULLIVAN 2015, PORTUGAL 2015, KNOX 2017 | **CITOKINES** | ↑ | Ifng |

**Table S2.**
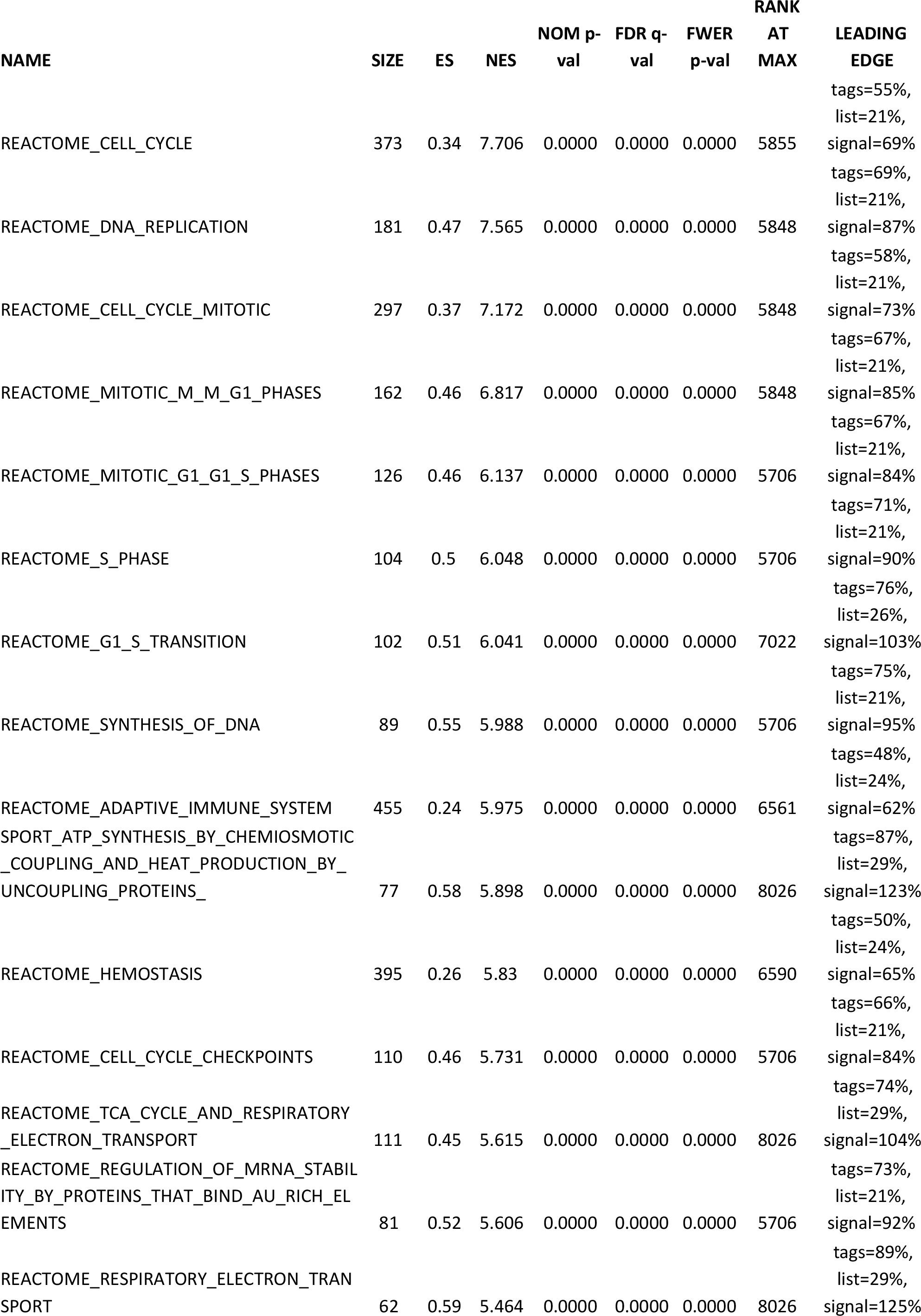
List of top 50 gene sets yielding the highest NES^a^ by GSEA.

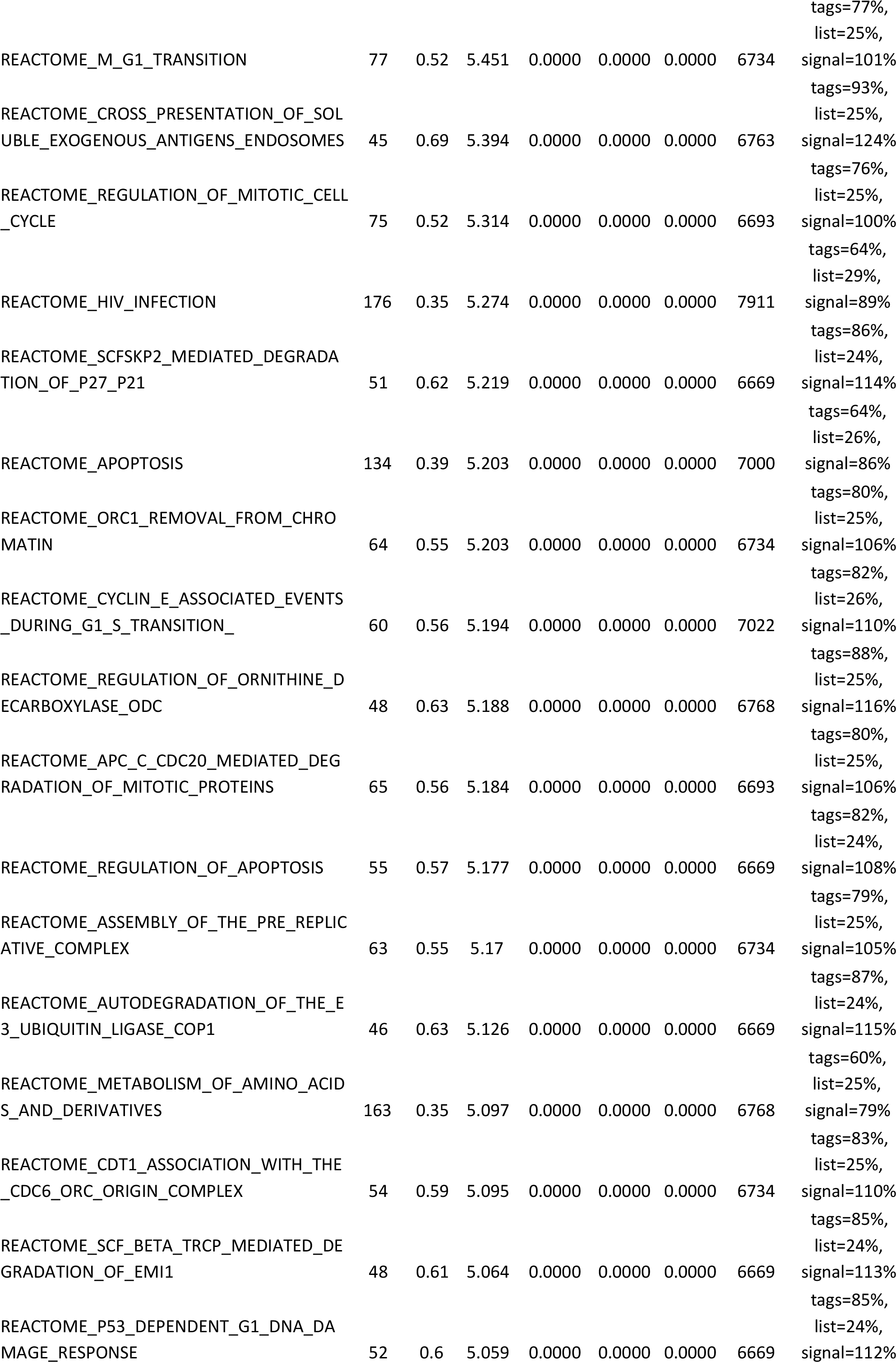

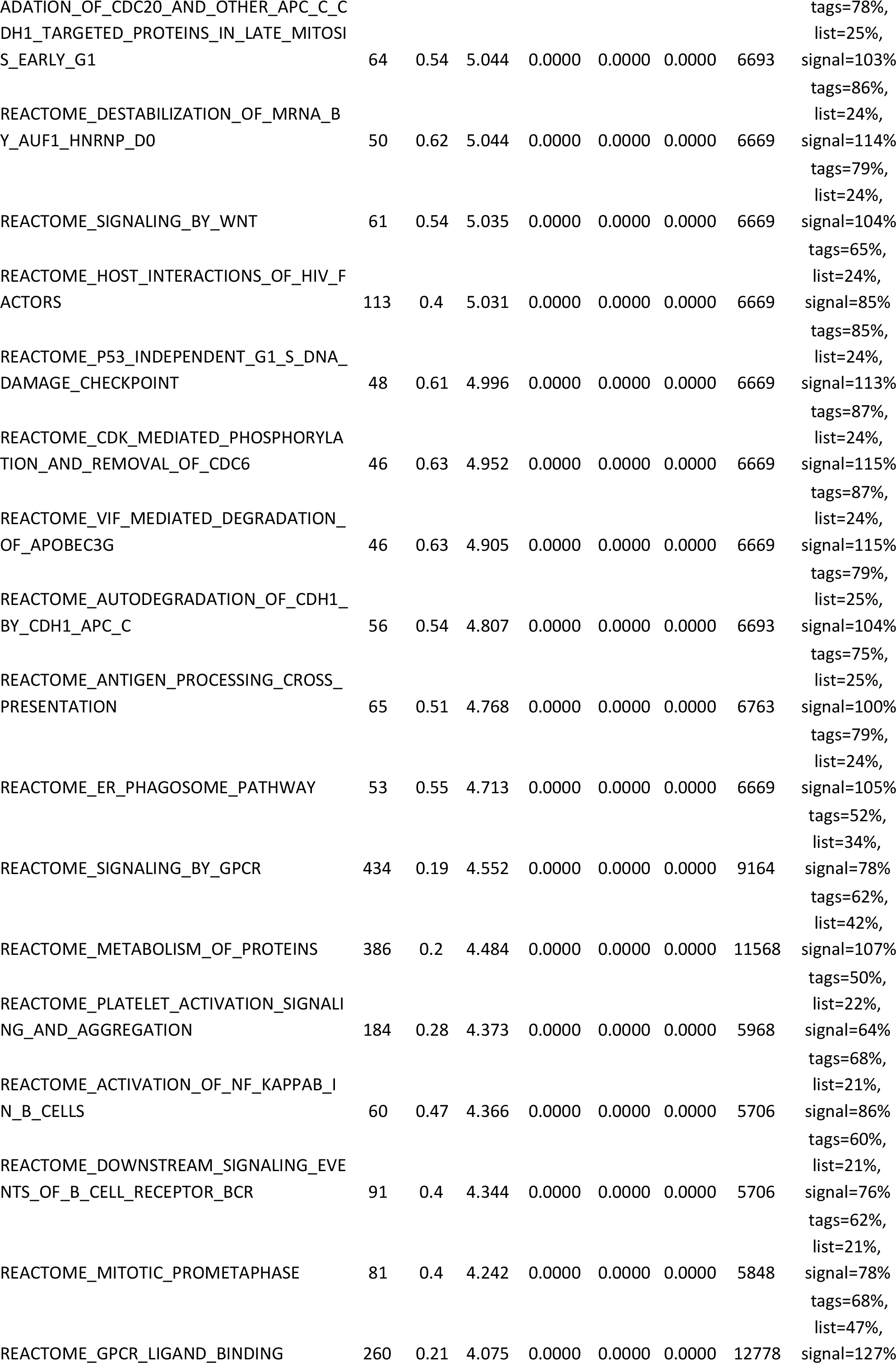

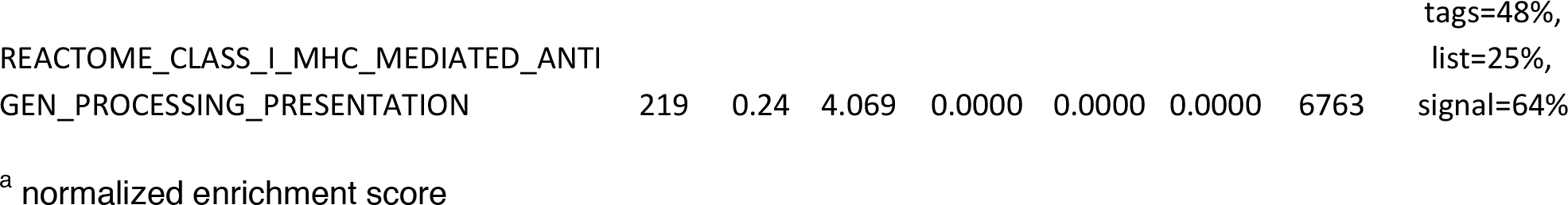

**Table S3.**
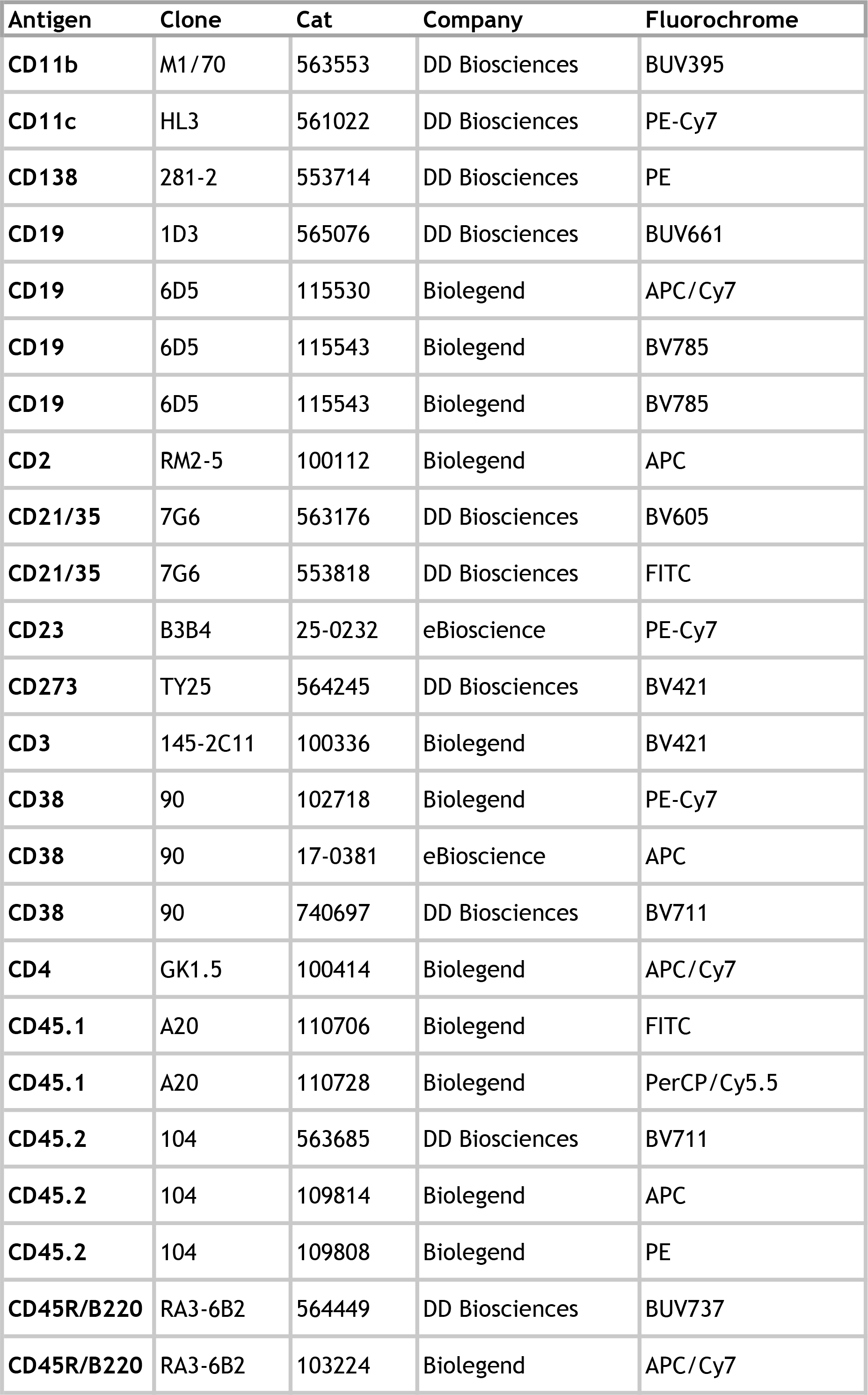
Antibodies used for flow cytometry.

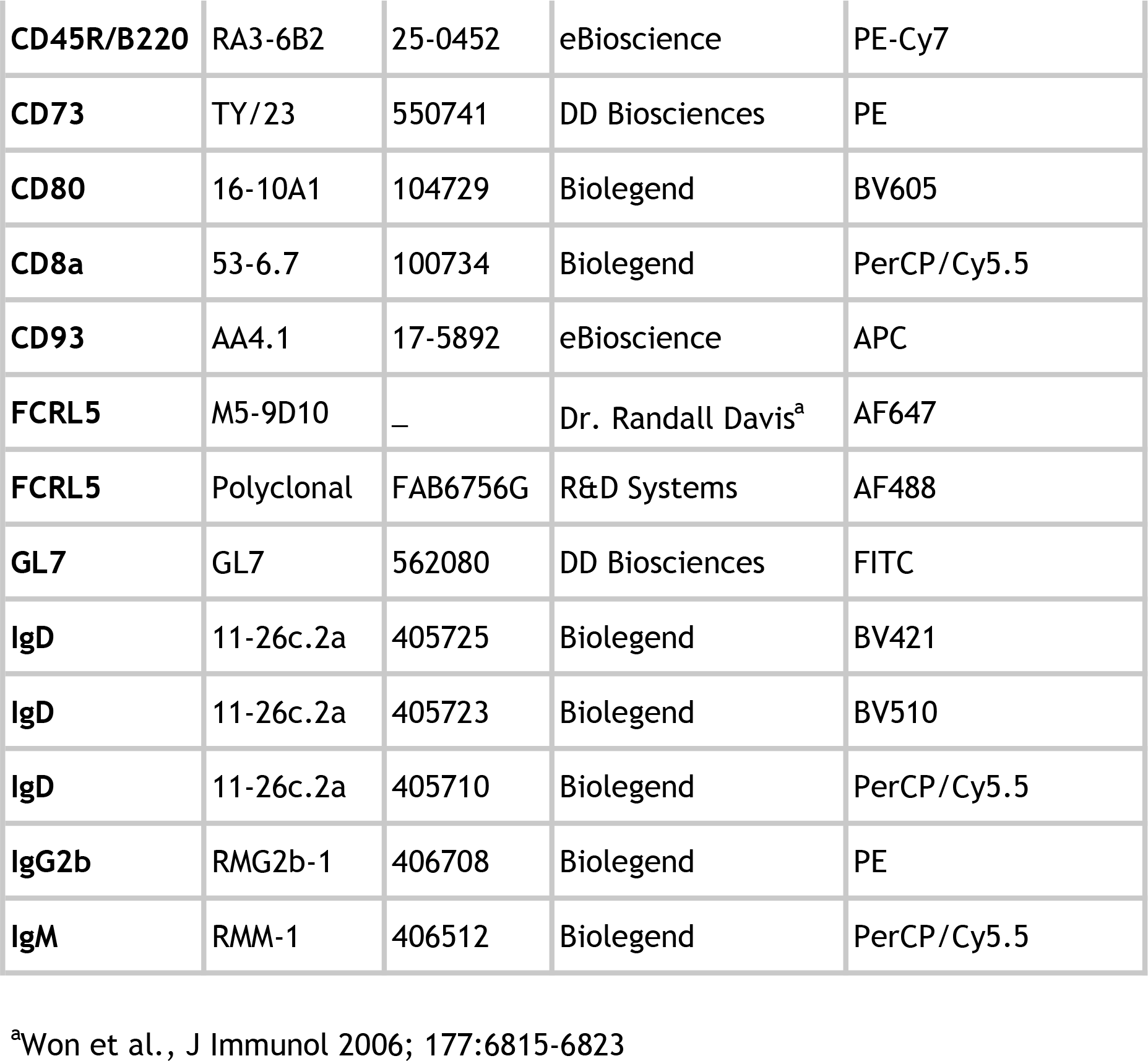

